# Chromosomal resolution reveals symbiotic virus colonization of parasitic wasp genomes

**DOI:** 10.1101/2020.03.19.994459

**Authors:** Jérémy Gauthier, Hélène Boulain, Joke J.F.A. van Vugt, Lyam Baudry, Emma Persyn, Jean-Marc Aury, Benjamin Noel, Anthony Bretaudeau, Fabrice Legeai, Sven Warris, Mohamed Amine Chebbi, Géraldine Dubreuil, Bernard Duvic, Natacha Kremer, Philippe Gayral, Karine Musset, Thibaut Josse, Diane Bigot, Christophe Bressac, Sébastien Moreau, Georges Periquet, Myriam Harry, Nicolas Montagné, Isabelle Boulogne, Mahnaz Sabeti-Azad, Martine Maïbèche, Thomas Chertemps, Frédérique Hilliou, David Siaussat, Joëlle Amselem, Isabelle Luyten, Claire Capdevielle-Dulac, Karine Labadie, Bruna Laís Merlin, Valérie Barbe, Jetske G. de Boer, Martial Marbouty, Fernando Luis Cônsoli, Stéphane Dupas, Aurélie Hua Van, Gaëlle Le Goff, Annie Bézier, Emmanuelle Jacquin-Joly, James B. Whitfield, Louise E.M. Vet, Hans M. Smid, Laure Kaiser-Arnault, Romain Koszul, Elisabeth Huguet, Elisabeth A. Herniou, Jean-Michel Drezen

## Abstract

Most endogenous viruses, an important proportion of eukaryote genomes, are doomed to slowly decay. Little is known, however, on how they evolve when they confer a benefit to their host. Bracoviruses are essential for the parasitism success of parasitoid wasps, whose genomes they integrated ~103 million years ago. Here we show, from the assembly of a parasitoid wasp genome, for the first time at a chromosomal scale, that symbiotic bracovirus genes spread to and colonized all the chromosomes. Moreover, large viral clusters are stably maintained suggesting strong evolutionary constraints. Genomic comparison with another wasps revealed that this organization was already established ~53 mya. Transcriptomic analyses highlight temporal synchronization of viral gene expression, leading to particle production. Immune genes are not induced, however, indicating the virus is not perceived as foreign by the wasp. This recognition suggests that no conflicts remain between symbiotic partners when benefits to them converge.

## Main

*Cotesia* wasps (Hymenoptera, Braconidae) are parasitoids of Lepidoptera. Female wasps lay their eggs into caterpillars and larvae develop feeding on the host hemolymph. Several *Cotesia* species are famous for their use as biological control agents to control insect pests, such as *Cotesia flavipes* which is massively released over several million hectares of sugarcane fields in Brazil^1,2^. Parasitoid wasps evolved several strategies that increase parasitic success, including a sensitive olfactory apparatus to locate their hosts^3,4^ and detoxification mechanisms against plant toxic compounds accumulating in their host (Fig. 1). But the most original strategy is the domestication of a bracovirus (BV) shared by over 46,000 braconid wasp species^5^. Bracoviruses originate from a single integration event ~103 million years ago (mya) of a nudivirus in the genome of the last common ancestor of this group^6–9^. Virus domestication nowadays confers a benefit to the wasps that use BVs as virulence gene delivery systems^10^. Indeed, virulence genes are introduced with wasp eggs into their hosts, causing inhibition of host immune defenses^10–12^. Imbedded in wasp DNA, the nudivirus genome has been extensively rearranged^13^. Now BV sequences are differentiated as i.) genes involved in particles production named “nudiviral genes”^6^, and ii.) proviral segments packaged as dsDNA circles in viral particles^13^, encoding virulence genes involved in successful parasitism.

**Fig. 1.**
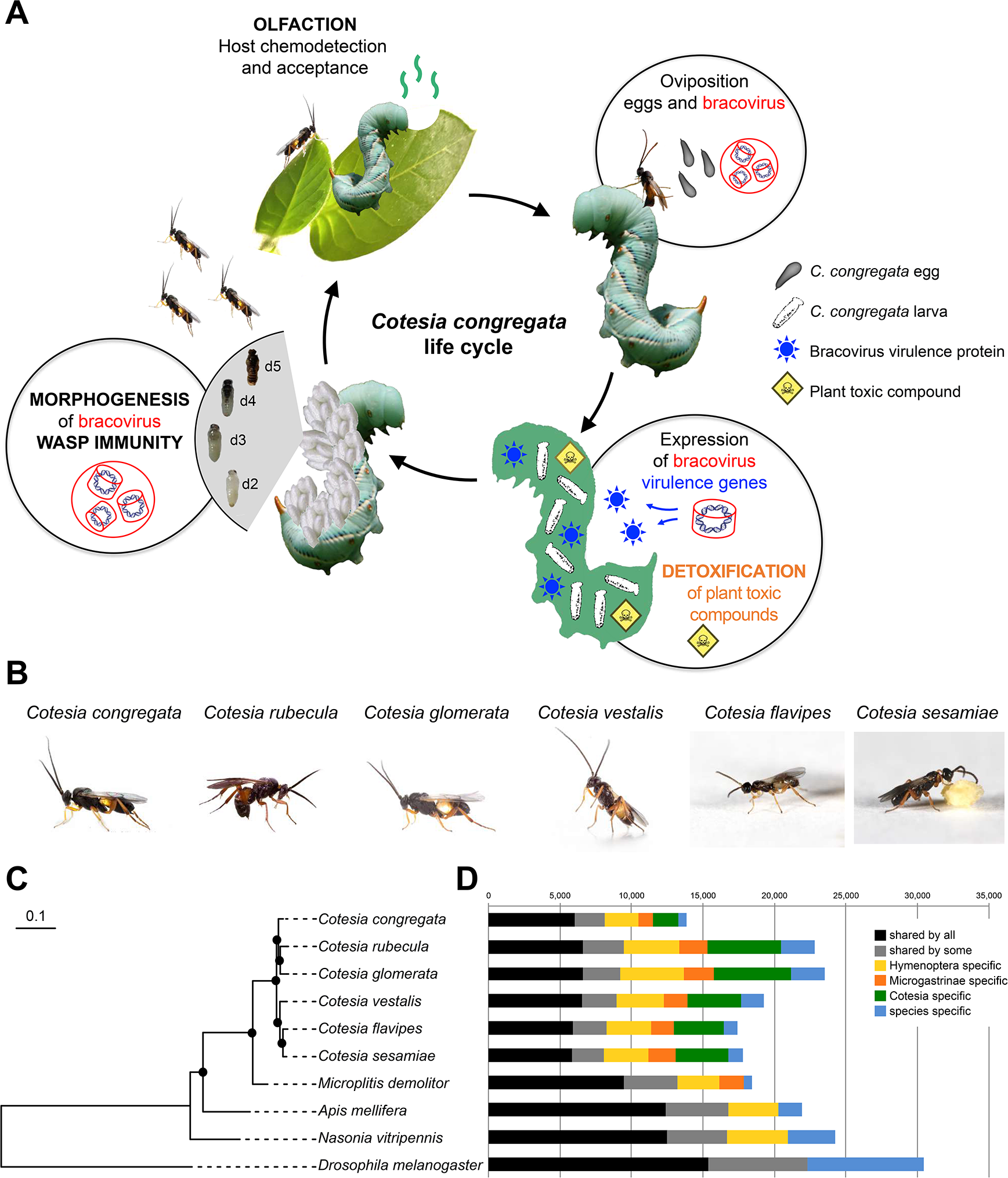
**A.** Major traits involved in the parasitoid koinobiont lifestyle and genome content of six *Cotesia* species. First OLFACTION plays an important role in the detection of the plant (tobacco) attacked by caterpillars and host (*M. sexta)* larvae acceptance by adult wasps. Once the host is accepted, the wasp injects its eggs bathed in ovarian fluid filled with bracovirus particles. Bracovirus particles infect host cells, from which expression of bracovirus VIRULENCE GENES alter host immune defenses, allowing wasp larvae development (the eggs laid in the host body would otherwise be engulfed in a cellular sheath of hemocytes). As the host ingests plant toxic compounds, such as nicotine, while feeding, wasp larvae consuming the hemolymph containing these compounds rely on DETOXIFICATION to complete their life cycle. But in these species associated with endogenous viruses the most important trait for parasitism success consists in BRACOVIRUS MORPHOGENESIS during wasp metamorphosis, using genes originating from a nudivirus ancestrally integrated in the wasp genome. As massive production of virus particles occurs within wasp ovaries, WASP IMMUNITY may be induced during particles production; d2, d3, d4, d5 refer to developmental stages of *C. congregata* larvae84. **B** Pictures of the six *Cotesia* species sequenced (credit H. M. Smid and R. Copeland). **C** Phylogeny of these species based on 1,058 single-copy orthologous insect genes including the Microgastrinae *Microplitis demolitor* and outgroups (*N. vitripennis*, *A. mellifera*, and *D. melanogaster*). Black dots highlight branches with at least 90% support from maximum-likelihood analysis (1,000 bootstraps). **D** Distribution of shared genes at several phylogenetic levels. Full protein-coding gene sets were included to identify orthologous gene groups. The “shared by some” category refers to genes shared by at least nine species among the 10 studied. Note that the lower number of genes for *C. congregata* probably reflects the higher quality of the genome assembly obtained.

To understand the genetic underpinning of the endoparasitoid life-style and of BV evolution since the original nudivirus capture event, we used a hybrid sequencing approach combining 454 reads, illumina short reads, and chromosomal contact data (HiC), to obtain a reference genome for *Cotesia congregata* at a chromosomal scale. These approaches allowed the analysis of wasp genes potentially involved in the success of the endoparasitoid lifestyle such as genes implicated in olfaction, detoxification and immunity. Moreover, the results revealed the global distribution of bracoviral sequences in wasp chromosomes and the identification of syntenies with genome scaffolds of *Microplitis demolitor* parasitoid wasp^14,15^, which diverged from the *Cotesia* lineage 53 mya^16^. In addition, we sequenced draft genomes of five *Cotesia* species (*C. rubecula*, *C. glomerata*, *C. vestalis*, *C. flavipes*, *C. sesamiae*) to assess bracovirus molecular evolution between closely related species.

We discovered that, in discrepancy with the usual gene decay of endogenous viral elements, bracovirus genes have spread in all wasp chromosomes, reaching a ~2.5-fold higher number of genes than a pathogenic nudivirus. However strong evolutionary constraints have maintained specialized chromosomal regions containing large virus gene clusters, resembling “super gene” regions and concentrating 77% of virulence genes and half of the nudiviral genes, respectively. Transcriptome analyses point to the major role of viral transcription in maintaining the viral entity, however wasp immune response is not induced, suggesting no conflicts remain in this ancient virus-wasp association.

## Results

### Genome assembly, annotation and comparison

First, a 207 Mb high-quality genome (scaffolds N50=1.1 Mb and N90=65 kb) was obtained for *Cotesia congregata* using a combination of mate pair 454 pyrosequencing and Illumina sequencing (Supplementary Table 1). Most of the assembly (86%) consisted of 285 scaffolds of over 100 kb. This genome was then re-assembled based on experimentally obtained chromosomal contact maps. The HiC method yielded ten main scaffolds comprising >99% of the previously obtained genome assembly (Supplementary Table 1 and 3; Supplementary Fig. 1 and 2), and corresponding to the ten chromosomes of *C. congregata*^17^. In addition, draft genomes of five related *Cotesia* species – *C. rubecula*, *C. glomerata*, *C. vestalis*, *C. flavipes* and *C. sesamiae* – were sequenced and assembled with Illumina shotgun sequencing reads (Supplementary Table 1). They respectively resulted in scaffold N50 values of 13, 9, 15, 20 and 26 Kb and cumulative sizes of 216, 243, 176, 155 and 166 Mb, in agreement with cytometry data (189-298 Mb) (Supplementary Table 2).

The genome of *C. congregata* comprised 48.7% of repeated DNA sequences including 34.7 % of known Transposable Elements (TEs) (Supplementary Fig. 3). Bracovirus proviral segments comprised other TE sequences that had previously been annotated as bracovirus genes: BV26 gene family corresponded to Sola2 elements (Supplementary Fig. 4; Supplementary Table 5) and CcBV 30.5 to a *hAT*^18^. Moreover, the analysis suggested that the BV7 gene family might also derive from TEs. Interestingly, some of these genes are highly expressed in the host^19^ or likely participate in ecological adaptation^20^. This indicates that, contrary to a common paradigm in this field^21^, the virulence genes packaged in bracovirus particles do not exclusively originate from the wasp cellular gene repertoire.

We annotated 14,140 genes in the genome of *C. congregata* (Methods and Supplementary Table 7), which include more than 99% of 1,658 conserved insect genes (98 to 99 % of the genes for the other *Cotesia* species, Supplementary Fig. 5).

### Gene gains and losses in traits associated to the endoparasitoid lifestyle

Expert annotation of chemoreceptor gene families identified 243 odorant receptor (OR), 54 gustatory receptor (GR) and 105 ionotropic receptor (IR) genes in *C. congregata*. These numbers are in line with those of other parasitoid wasps, but are lower than in ants22 (Supplementary Table 10). Phylogenetic analyses showed *C. congregata* ORs belong to 15 of the 18 OR lineages (Fig. 2) described in Apocrita^23^ and revealed independent OR gene expansions in *N. vitripennis* and in Braconidae (Fig. 2). The most spectacular Braconidae-specific expansions occurred in five clades each harboring at least 25 genes in *C. congregata* (Fig. 2; Supplementary Table 10). Highly duplicated OR genes were found in 6 clusters of at least 10, and up to 19, tandemly arrayed genes (Supplementary Fig. 16). Within Braconidae, many duplications occurred in the ancestors of *Cotesia* but OR copy numbers varied significantly between species (Fig. 2). This illustrates the highly dynamic evolution of OR gene families within parasitoid wasps and between *Cotesia* species, which have different host ranges.

**Fig. 2.**
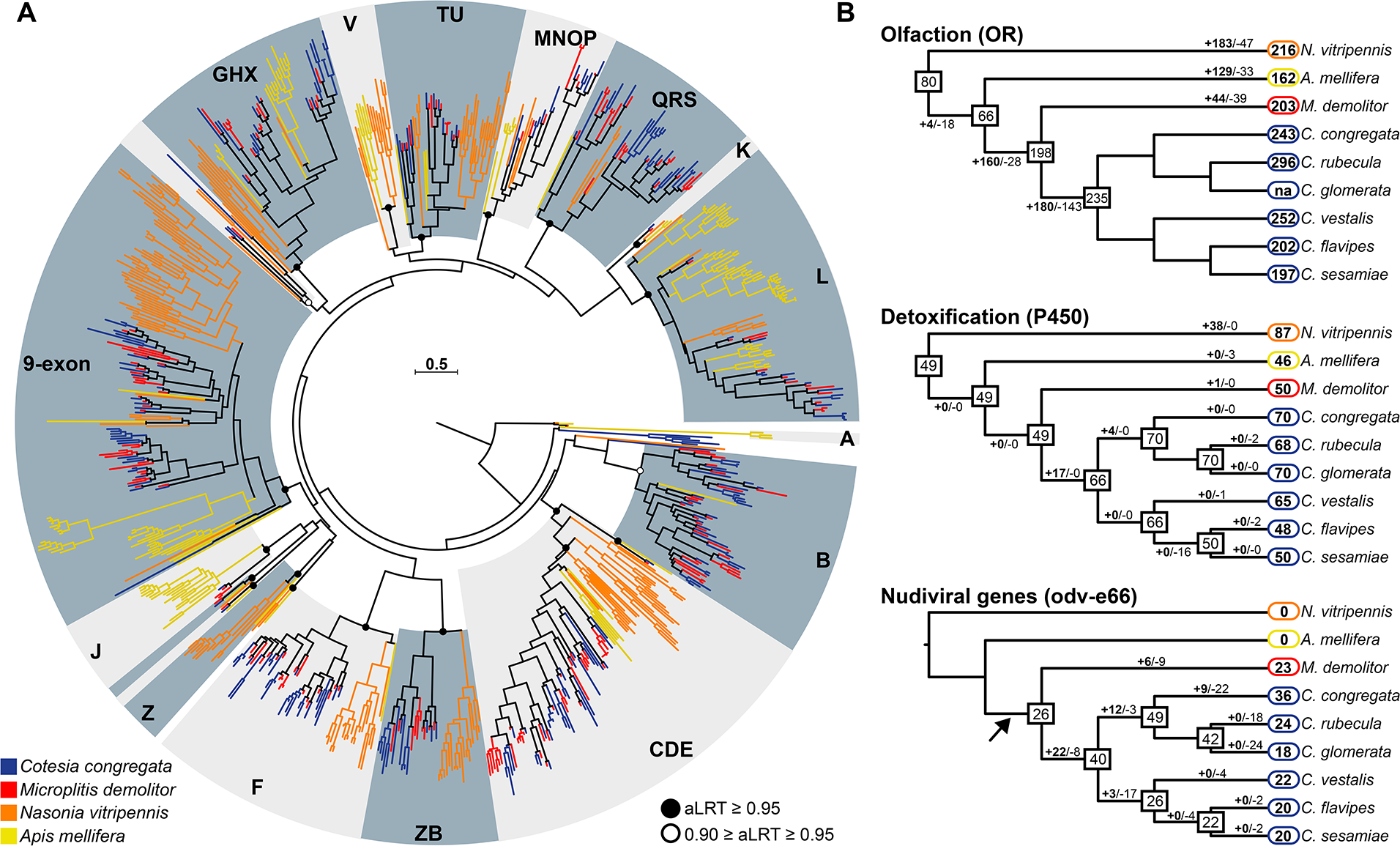
Gene family extensions in *Cotesia.* **A** Maximum-likelihood phylogeny of the OR family in *C. congregata* and four other Hymenoptera species. The dataset included 243 amino acid sequences from*C. congregata* (blue), 203 sequences from *M. demolitor* (red), 216 sequences from *N. vitripennis* (orange), 162 sequences from *A. mellifera* (yellow). The tree was rooted using the Orco (OR-coreceptor) clade. Circles indicate nodes strongly supported by the approximate likelihood-ratio test (black circles aLRT ≥ 0.95; white circles 0.90 ≥ aLRT ≤ 0.95). The scale bar represents 0.5 expected amino acid substitutions per site. ORs of the five Hymenoptera species are distributed into 18 OR subfamilies previously described in 23 delineated in grey. **B** Copy number dynamics of OR (olfaction) P450 (detoxification) and Odv-e66 genes, note that the later are found specifically in bracovirus associated wasps since they derive from the ancestrally integrated nudivirus. Estimated numbers of gene gain and loss events are shown on each branch of the species tree. The size of OR repertoires in common ancestors is indicated in the boxes. The lack of phylogenetic resolution for closely related Cotesia OR genes precluded any comprehensive analysis of gene gains and losses.

All the genes involved in detoxification in insects were identified in *C. congregata* and are largely conserved within *Cotesia* (Supplementary Table 12 and 13). For instance, each species harbors conserved numbers of UDP-glucosyltransferases (UGTs) and slightly different numbers of gluthatione-S-transferases (GSTs). In contrast, carboxylesterases (CCEs) and cytochrome P450 (P450s) numbers vary widely with *C. flavipes* and *C. sesamiae* harboring few representatives (respectively 22-24 CCEs and 49 P450s), compared to the 32 CCEs of *C. rubecula* and the 70 P450s found in *C. congregata*, which are both exposed to plant toxic compounds (Supplementary Table 12 and 13). *Cotesia* specific P450 families were identified in the clan 3 and 4, both of which are often associated to adaptation to plant compounds and insecticides^24^ (Supplementary Table 12 and 13). Altogether *Cotesia* appear equipped for detoxification although no spectacular gene expansion was observed.

The complete genome annotation of *C. congregata* revealed 102 nudiviral gene copies that have colonized all the chromosomes. This number is similar to that of pathogenic nudiviruses^25^, which is an unexpected result given that endogenous viral elements usually undergo gene loss in the course of evolution^26^. Here this surprisingly high number of nudiviral genes can be explained by the fact that gene losses have been balanced by expansions of gene families. At least 25 of the 32 nudivirus core genes involved in essential viral functions^27^ have been retained in the wasp genome, with the notable absence of the nudiviral DNA polymerase (Supplementary Table 8). The loss of this major replication gene may have prevented viral escape^10,25,28^. The *fen* genes, generally involved in DNA replication, form a gene family with six tandem copies that is found specifically in the *Cotesia* lineage (Fig. 3B). The most spectacular expansion concerns the *odv-e66* gene family with 36 genes in 10 locations (Fig. 2B; Fig. 3C), in contrast to the one or two copies typically found in nudivirus genomes^29^. This expansion occurred both before and after the divergence between *C. congregata* and *M. demolitor* since we found tandemly duplicated copies in homologous loci of the two species or in *C. congregata* only (Fig. 3D). In baculoviruses, *odv-e66* encodes a viral chondroitinase^30^ involved in digesting the peritrophic membrane lining the gut, thus allowing access to target cells during primary infection. Different ODV-E66 proteins may similarly allow BVs to cross various host barriers, and BV infection to spread to virtually all Lepidoptera tissues^31,32^.

**Fig. 3.**
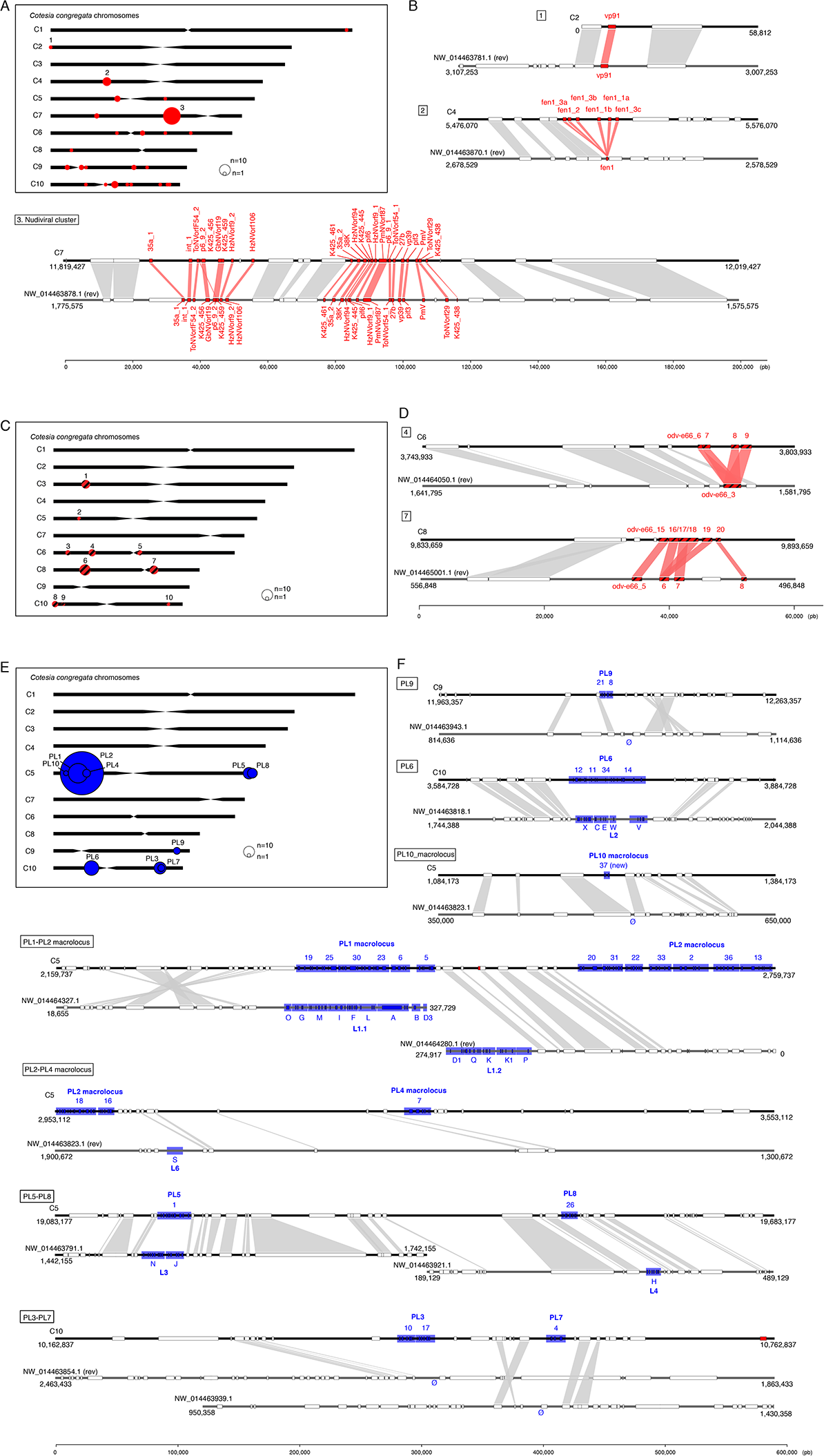
Synteny of nudiviral genes loci and Proviral Loci (PL) between *C. congregata* and *M. demolitor*. **A** *C. congregata* chromosome map with the position of 24 nudiviral genes loci. **B** Comparisons between nudiviral gene regions of *C. congregata* and *M. demolitor*. Synteny between the two species has been characterized by at least two hymenopteran (non-viral) orthologous genes in the vicinity of homologous nudiviral gene(s) of both species. Genome scaffolds are represented in black. Red boxes indicate nudiviral genes and white boxes refer to hymenopteran genes. 1. the *vp91* region is orthologous indicating the position of this gene was inherited from their common ancestor 53 mya; 2.: the fen region is also orthologous but an expansion occurred specifically in *Cotesia* lineage giving rise to six copies; 3.: the organization of the nudiviral cluster encoding in particular capsid genes has remained strikingly similar with the same viral genes in the same order (except p6-9-2) in both species indicating strong evolutionary constraints. **C** *C. congregata* chromosome map with the position of gene loci corresponding to the highly expanded *odv-e66* nudiviral gene family. **D** Comparison of two odv-e66 loci showing that expansion occurred before (cluster 7) and after (cluster 4) the separation of both species **E** *C. congregata* chromosome map with the position of Proviral Loci (PL) encoding virulence genes packaged in bracovirus particles. Note the concentration of loci (successively PL10-PL1-PL2 and PL4) in a 2 Mb region termed “macrolocus” and representing half of the chromosome 5 short arm. **F** Comparison of *C. congregata* and *M. demolitor* PL. Numbers 1 to 37 and letters correspond to the different dsDNA circles present in CcBV and MdBV particles produced from the PL. Blue boxes indicate virulence genes while white boxes refer to hymenopteran genes and the red box to a nudiviral *odv-e66* gene located between PL1 and PL2. Ø indicates the absence of orthologs PL in the *M. demolitor* genome.

### Genomic architecture and synteny of bracovirus genes

Chromosome scale genome assembly of *C. congregata* provides the first comprehensive genomic organization of a bracovirus within the genome of a wasp. Half of the single copy nudiviral genes are located in the nudiviral cluster, which comprises 25 genes and is amplified during bracovirus production but not packaged within the particles^33^. Comparison with the scaffolds of *M. demolitor* showed overall gene content conservation as well as conserved syntenic blocks in the genomic regions flanking nudiviral sequences, over ~53 million years of evolution. This confirms that the nudiviral clusters of both species are orthologous (Fig. 3) and likely derive from a genome fragment of the nudivirus that originally integrated in the ancestor wasp genome. This striking stability suggests that major evolutionary constraints maintain these genes together. The other nudiviral genes are dispersed in the wasp genome, although not evenly, as more loci are located in the 4 smaller chromosomes. Orthology with *M. demolitor* could be identified for 20 nudiviral gene regions (Fig. 3; Supplementary Fig. 9), indicating they were already dispersed in the last common ancestor of both wasps and have stayed in the same loci. Altogether, this showed that nudivirus gene loss and dispersion occurred during the first period of wasp-bracovirus association (100 to 53 mya).

The expansion of virulence genes is another aspect of wasp genome colonization. In *C. congregata,* the 249 virulence genes encoded by proviral segments are found in 5 regions of the genome located on three chromosomes. However, 77% of these genes clustered in a single region, which comprised four physically linked proviral loci (PL1, PL2, PL4, PL10) interspersed by wasp genes (Fig. 3). This major virulence gene coding region, named “macrolocus”^13^, spans half of chromosome 5 short arm and is almost comparable (2 Mb, 177 genes) to the Major Histocompatibility Complex (MHC) involved in mammalian immunity (~4 Mb, ~260 genes)^34^. Orthology relationships could be inferred between the PL1 in *C. congregate* macrolocus and the largest proviral region (comprising 11 segments) of *M. demolitor*^14^ (Fig. 3) but the macrolocus expanded specifically in the *Cotesia* lineage since *M. demolitor* does not have regions corresponding to PL2, PL4 and PL10. Further syntenies were found between 5 isolated proviral loci, showing they pre-date wasp lineage split (Fig. 3) whereas the 3 proviral loci in the long arm of chromosome C9 and C10 appear as a genuine novelty of the *Cotesia* lineage. The homologous relationships of PLs contrasts with their gene content divergence^35^, which show high turnover probably because of their role as virulence factors and coevolution with the hosts^20^.

### Strong conservation of the bracoviral machinery

The DNA circles packaged in bracovirus particles are produced following the genomic amplification of replication units (RU) that span several proviral segments of PLs^33,36^. Detailed genomic analyses of *C. congregata* data led to the identification of a specific sequence motif at each RU extremity and confirmed the presence of circularization motifs^13,37^ on all proviral segments at the origin of packaged circles indicating the conservation of the viral machinery whatever the localization of viral sequences. (Fig. 4; Supplementary Fig. 10).

**Fig. 4.**
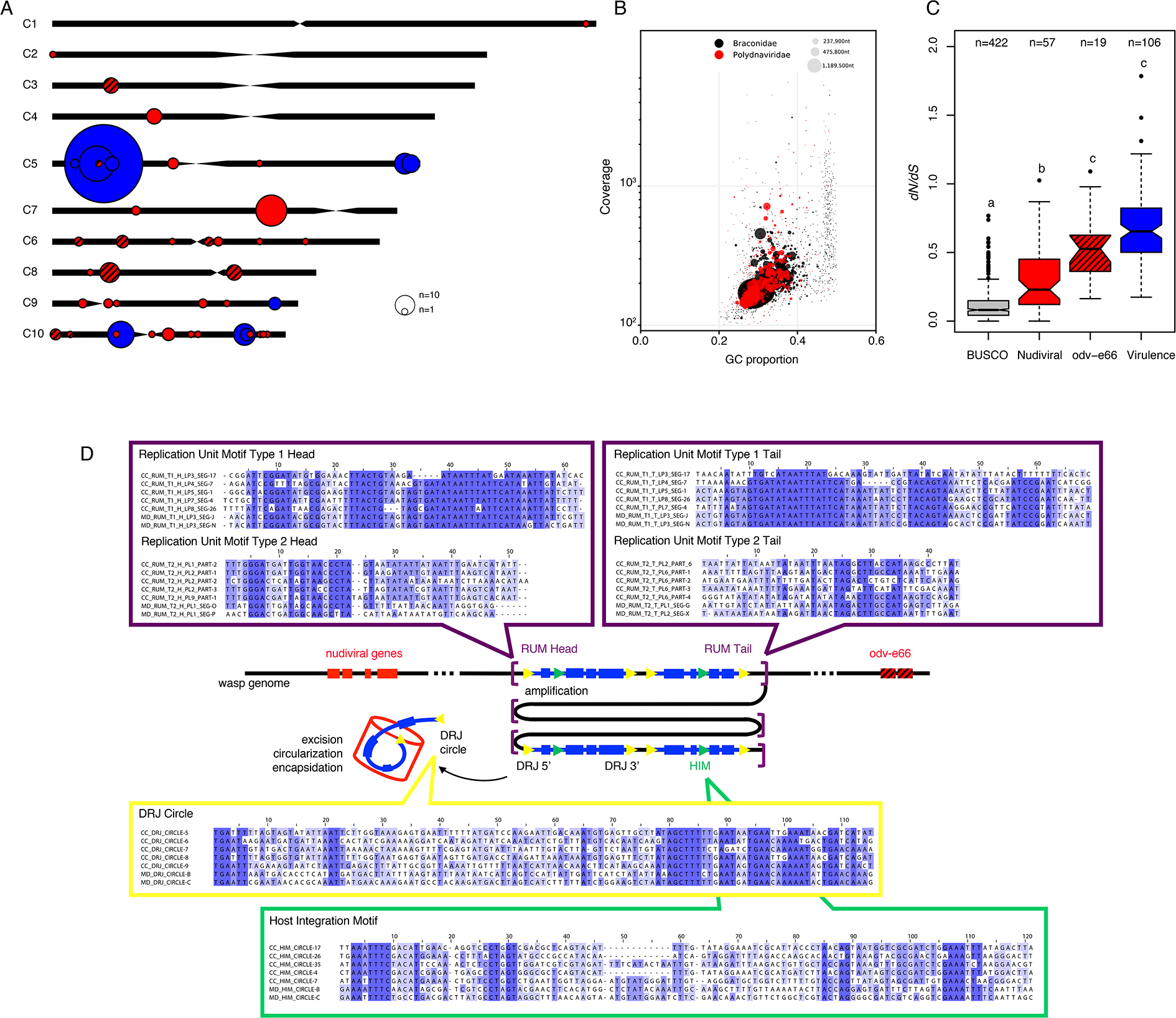
**A** *C. congregata* chromosome map with the location of bracovirus loci: nudiviral gene loci are shown in red, nudiviral *odv-e66* gene loci in hatched red and Proviral Loci (PL) in blue. The sizes of the circles correspond to the relative number of genes in each locus **B** taxon-annotated GC content-coverage plot of *C. congregata* scaffolds. Each circle represents a scaffold in the assembly, scaled by length, and colored by taxonomy assignation to Braconidae and Polydnaviridae. The x-axis corresponds to the average GC content of each scaffold and the y-axis corresponds to the average coverage based on alignment of illumina reads. **C** Measure of selection pressure on hymenopteran conserved genes, nudiviral genes and virulence genes. Pairwise evolutionary rates (dN/dS) of single-copy orthologous BUSCO genes, nudiviral genes, different copies from the expanded odv-e66 nudiviral gene family and virulence genes of *C. congregata* and *C. sesamiae*. Letters above boxes indicate significant differences determined by Kruskal–Wallis test (H= 296.8, 2 d.f., P<0.001) followed by *post-hoc* comparisons. **C** Schematic representation of the genomic amplification during the production of viral particles in the wasp ovaries. Replication Unit Motifs (RUM) are the motifs that constitute the extremities of the molecules amplified during particle production. Direct Repeat Junctions (DRJ), at the extremities of each segment are used during the excision/circularization process to produce packaged dsDNA circles from the amplified molecules. Host Integration Motifs (HIM) are motifs used during the integration of bracovirus circles in host genome. For each of these motifs an alignment of a representative set of sequence comprising five motifs from *C. congregata* and *M. demolitor* are represented (all alignments are reported in Supplementary Fig. 10).

Over 100 million years of evolution, the conservation of viral functions in microgastroid wasps must have endured through several bouts of natural selection. Synonymous to nonsynonymous substitution ratio analyses on orthologous nudiviral genes in *Cotesia* showed most of the nudiviral genes (65 genes among the 79 tested genes), are evolving under conservative selection, that is however less stringent than on the insect genes used to assess genome completeness (Fig. 4; Supplementary Table 8). This selection is notably strong for genes involved in viral transcription, such as the RNA polymerase subunits (*lef-4*, *lef-8*, *lef-9*, *p47*), which most likely control nudiviral gene expression and, consequently, particle production^6,8^. In contrast, genes involved in infectivity (homologues of baculovirus *pif* genes) appear less conserved (Supplementary Table 8). This might reflect divergence occurring during host shifts, through adaptation of virus envelope proteins to particular host cell receptors. The large *odv-e66* gene family and duplicated genes (*p6.9_2*, *pif-5_2, 17a*) similarly displayed less stringent to relaxed selection (Fig. 4; Supplementary Table 8), which might be conducive to mutational space exploration for adaptation by neo-functionalization or sub-functionalization38. Virulence genes encoded by proviral segments globally displayed lower conservative selection than nudiviral genes (Fig. 4), as expected for genes interacting with host defenses and involved in evolutionary arms race or adaptation to new hosts^20^.

### Synchronized nudiviral transcription precedes bracovirus production

Bracovirus particle production only occurs in the calyx region of the ovaries, and starts before wasp emergence at day 5. Detailed RNAseq analysis of wasp ovaries and venom glands during pupal development showed 91 (out of 102) nudiviral genes were only expressed in the ovaries and could not be detected in the venom gland, which highlights strong ovarian tissue specificity. The onset of nudiviral gene expression was detected from day 2 of pupal development, with the expression of the RNA polymerase subunits (Fig. 5), reaching a peak on day 4 and declining in later stages (Fig. 5). The other nudiviral genes followed roughly the same expression pattern, low or undetectable at day 2, high at day 3 and reaching a peak at day 5 (Fig. 5). This pattern concurs with the hypothesis that the nudiviral RNA polymerase controls the expression of the other nudiviral genes and therefore bracovirus particle production. Notably, 12 nudiviral genes reached very high mRNA levels (at day 5) and are among the top 50 of most expressed genes in *C. congregata*. Three genes from the nudiviral cluster (including a major capsid component vp39) are by far the 3 most expressed of all wasp genes. Altogether, it is remarkable that the expression of almost all nudiviral genes remains strongly synchronized during pupal ovarian development. Considering the age of the nudivirus wasp association, this suggests strong evolutionary constraints.

**Fig. 5.**
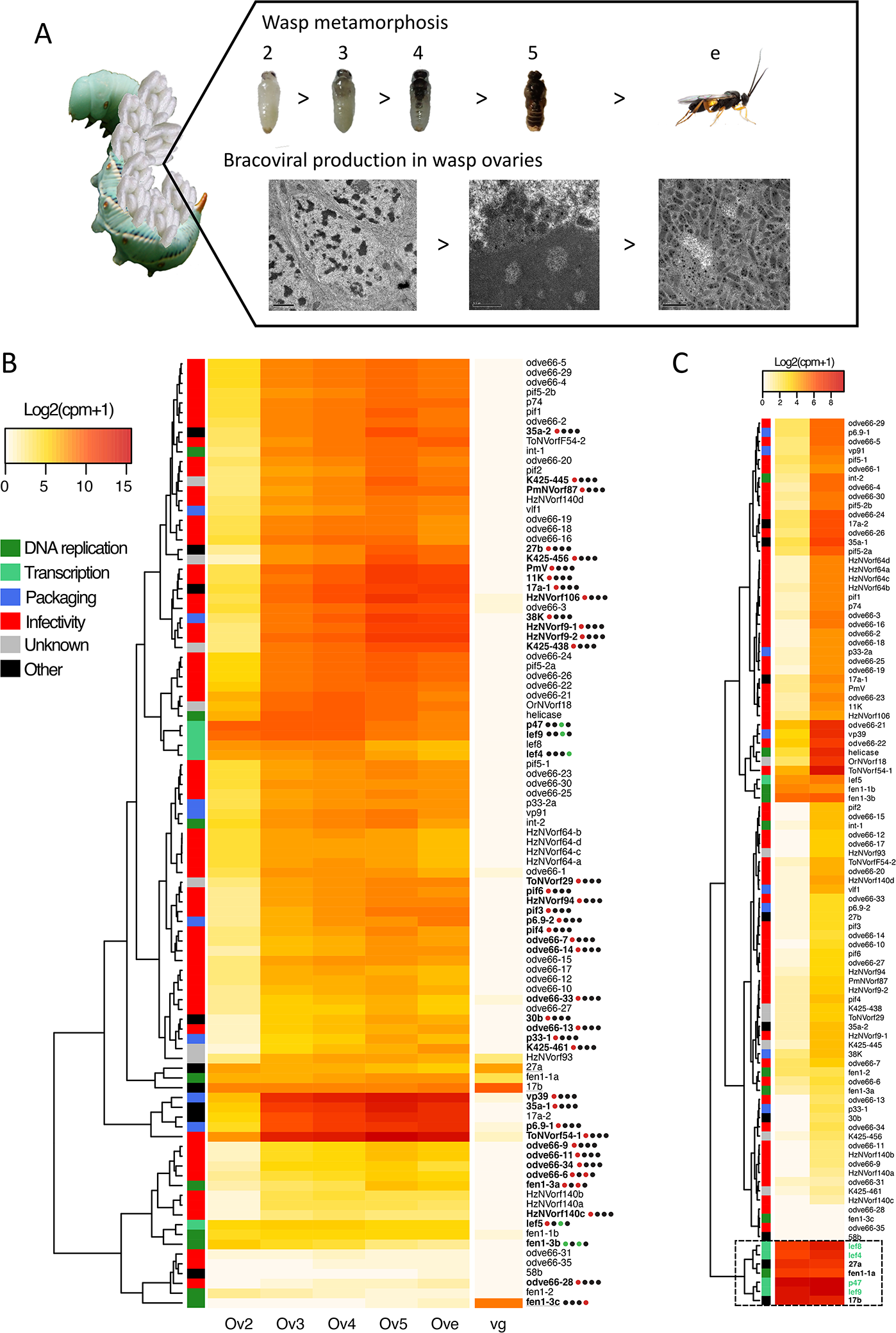
Gene expression of nudiviral genes in the ovaries during *C. congregata* nymphal development. **A** Pictures of wasp developmental stages studied and characteristic electron micrographs of ovarian cells involved in particle production. From d2 to d4 stage, cells that will produce particles show enlarged nuclei with chromatin condensation (left panel). From d5 massive particle production begins, particle assembly occurs at the periphery of a zone of electron dense material in the nucleus named “virogenic stroma” (middle panel). In newly emerged wasp nuclei are completely filled with bracovirus particles (right panel). Credit: Juline Herbinière. **B** Unsupervised hierarchical clustering based on gene expression in ovaries and venom glands. Ov2, Ov3, Ov4, Ov5 refer to the different ovaries across wasp developmental stages. Ove and vg refer to ovaries and venom glands of adult wasps. The colored squares associated with the clustering tree indicate the viral functions to which different nudiviral genes are supposed to contribute based on those of their baculovirus homologues. Heatmap of expression levels of 95 nudiviral genes is shown in the middle panel. Bold names highlight the genes that are validated as significantly differentially expressed between two consecutive stages using the statistical analysis and dots represent the four different comparisons studied between ovary stages (Ov2 *vs.* Ov3, Ov3 *vs*. Ov4, Ov4 *vs*. Ov5 and Ov5 *vs.* Ove). Black, red and green dots indicate similar, increased and reduced expressions between consecutive developmental stages respectively. The increase of some nudivirus genes expression between d2 and d3 visualized on the heat map was not validated statistically for all of them because in one of the d2 duplicates (shown in C) nudiviral genes expression had already reached high levels. Underlined genes show higher expression in venom glands compared to ovaries (Ove) note that 27a and 17b are not nudiviral genes but wasp genes, the products of which have been identified in Chelonus inanitus bracovirus particles. **C** Unsupervised hierarchical clustering of gene expression from the two replicates of Ov2 ovary stage, that are very different regarding nudiviral gene expression levels, although dissected nymphae presented a similar coloration pattern, the left one representing a slightly earlier stage from the analysis of the whole set of wasp genes. Note that the genes within the box are already expressed at a high level in the earlier stage, including all of the genes involved in nudiviral transcription (shown in green) except lef5, in accordance with the hypothesis that the nudiviral RNA polymerase complex controls the expression of the other genes (lef-5 is associated to the complex but not a part of it).

### Immune gene expression during bracovirus production

After 100 million years of endogenous evolution within the wasp genome, one can question whether the bracovirus is considered as a regular part of the wasp or is still perceived as non-self, in which case its production should trigger an immune response. However, whether viral production interacts in any way with the wasp immune system was totally unknown. Globally, the annotation of immune-related genes indicated *C. congregata* has an arsenal of 258 immune genes that are potentially involved in recognition, signal transduction, different signaling pathways, melanization and effector functions (Supplementary Table 9), in accordance with the recently reported annotation of *C. Vestalis* immune genes^39^. We identified all members of the Toll, IMD, Jak/STAT and RNA interference pathways found in Hymenoptera (Supplementary Table 9).

In contrast to the sharp increase in nudiviral gene expression, no significant changes in immune genes expression could be detected in the ovaries during pupal development (Fig. 6). In particular, expression of the genes involved in antiviral immunity (encoding members of the RNA interference, Jak/STAT or Imd/JNK pathways) was high in ovaries, even at stages before particle production is observable (ovaries stages 2 to 4), but hardly fluctuated during the course of ovary development, even at day 5, when massive particle production becomes apparent by TEM (Fig. 5A). Thus, no immune response appears to be induced or repressed at the cellular level as a response to high level nudiviral genes expression and particles production. This strongly suggests the bracovirus is not perceived as foreign by wasp ovarian cells (Fig. 1).

**Fig. 6.**
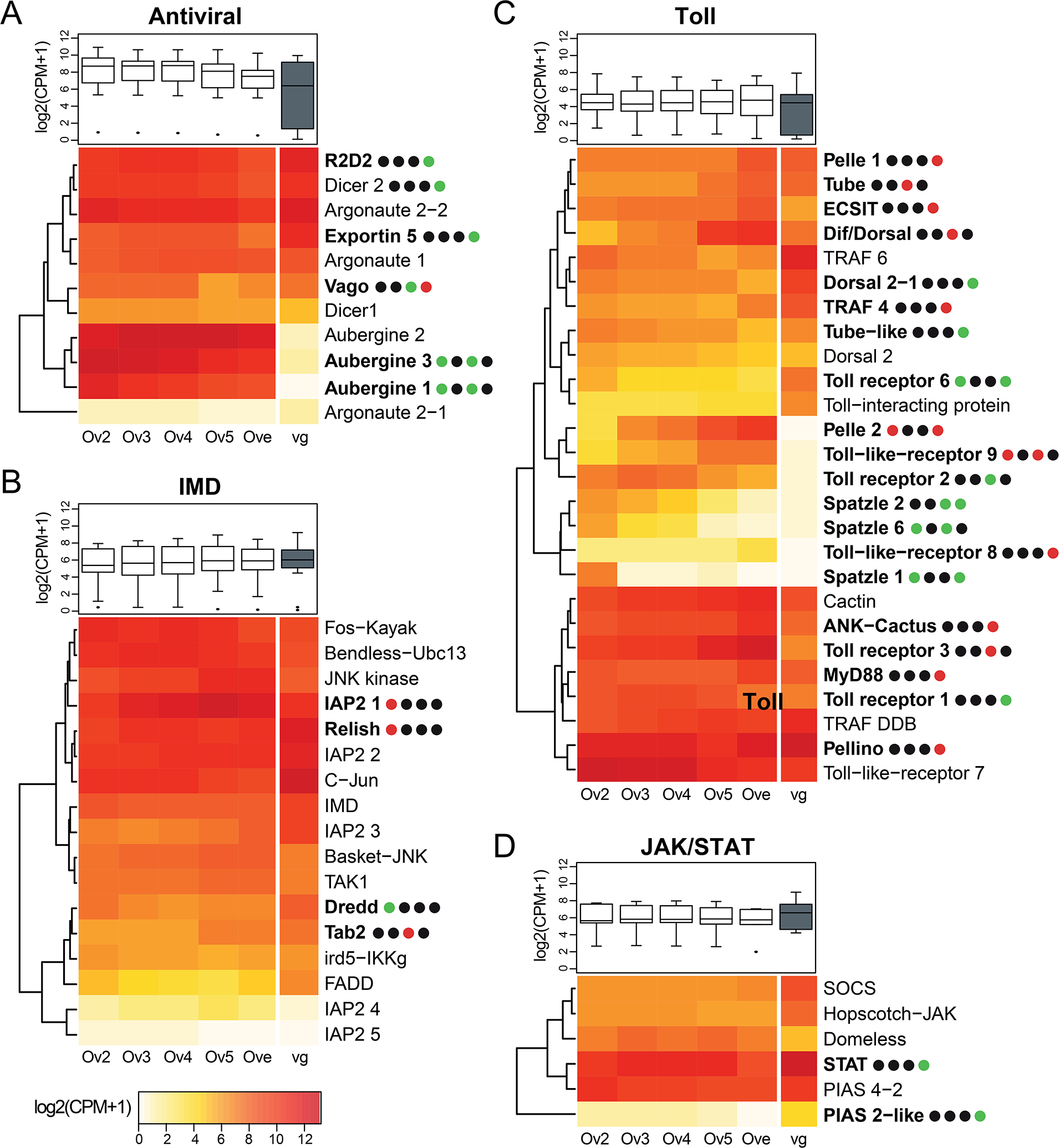
Gene expression of immune genes in the ovaries during *C. congregata* nymphal development. Gene expression of antiviral immunity genes during *C. congregata* development. The heatmaps show the expression of genes involved in **A** RNAi, **B** Imd, **C** Toll and **D** Jak-STAT pathways across the developmental stages of ovaries (Ov2, Ov3, Ov4, Ov5, Ove) and in venom glands (vg). The trees on the left are unsupervised hierarchical clustering of expression values. Boxplots represent overall expression of each pathway in ovaries and venom glands. Bold names highlight the genes that are differentially expressed between two stages and dots represent the four different comparisons studied between consecutive ovary stages (Ov2 *vs.* Ov3, Ov3 *vs*. Ov4, Ov4 *vs*. Ov5 and Ov5 *vs.* Ove). Black, red and green dots indicate similar, increased and reduced expressions between consecutive developmental stages respectively. Note that no particular trend appears correlated to bracovirus particles production which occurs massively from Ov5 onward.

## Discussion

To investigate the genome features related to the endoparasitoid lifestyle, we sequenced the genomes of 6 *Cotesia* species, and obtained an assembly at the chromosomal scale for *C. congregata.* This approach provided insights into essential functions of the wasps, such as olfaction, detoxification and immunity, as well as on the genomic evolution of the bracovirus. Large OR gene diversifications are often associated to host localization and acceptance. Indeed, female wasps rely on sensitive olfactory apparatus to locate their hosts from volatile cues plants emit in response to herbivore attacks^3,4^ and to assess caterpillar quality before oviposition^40^. Interestingly, OR copy numbers varied significantly during the evolution of the genus *Cotesia* (Fig. 2). The high dynamics of OR repertoire might point to the need for more specific recognition of chemical cues from the host and its food plant.

In contrast, no comparable diversification was observed in the detoxification arsenal, even though *Cotesia* larvae can be exposed to various toxic phytochemicals while feeding on the hemolymph of caterpillar hosts (i.e. potential exposure of *C. congregata* to insecticidal nicotine when parasitizing *Manduca sexta* feeding on tobacco; *C. rubecula*, *C. glomerata* and *C. vestalis* are potentially exposed to glucosinolates by developing in hosts consuming crucifers; and *C. flavipes* to phenylpropanoids and flavonoids when parasitizing hosts on sugarcane). This surprisingly low diversification of the detoxification arsenal could suggest that wasp larvae may not be as exposed to toxic compounds as expected due to direct excretion of these chemicals by the host larvae^41,42^.

*Cotesia* wasps face different immune challenges during their lifetime. While feeding on nectar the adult might be exposed to similar environmental challenges as honey bees. Development inside the caterpillar host could on the one hand shield wasp larvae from pathogens but on the other hand expose them to opportunistic infections, because parasitism alters caterpillar immune defenses. Lastly, metamorphosis coincides with the production of bracovirus particles, against which wasp antiviral responses had so far not been investigated. Insects were recently shown to recognize their obligate bacterial symbionts as foreign and to exert strong immune control, as documented for *Sitophilus oryzae* Primary Endosymbiont (SOPE)^43^. As the immunity gene arsenal of *C. congregata* is comparable to that of the honey bee, this wasp is most probably able to mount an immune response against pathogens, including viruses. However, the transcriptomic approach did not reveal any significant difference in immune gene expression between the ovaries of different pupal stages, although massive amounts of bracovirus particles are produced from day 5. This lack of anti-viral gene induction suggests the wasp no longer perceives the bracovirus as potentially pathogenic but rather considers it as a regular secretion.

With the ancestral integration of a nudivirus genome, the parasitoid wasp gained a series of viral functions: including viral transcription, viral DNA replication, particle morphogenesis and infectivity. Whereas the function of viral DNA replication has been lost^10^, thus impeding autonomous virus re-emergence, the other viral functions have been reorganized for virulence gene transfer via bracovirus particles. Chromosomal scale resolution of *C. congregata* genome showed bracovirus genes have colonized all the chromosomes with a nearly 2.5 fold increase of in the total number of virus genes compared to the genome of a pathogenic nudivirus. This contrasts sharply with the decay of most viruses integrated into eukaryote genomes that do not provide a benefit to their host. Bracovirus dispersion occurred between 100 and 53 mya, as 25 viral loci are orthologous between *Cotesia* and *Microplitis* (Fig. 3; Supplementary Fig. 5). Yet, the organization of many bracovirus genes in clusters suggests strong evolutionary constraints maintain these genes together. In the case of the nudiviral cluster, which encodes major capsid components (VP39, 38K) on chromosome 7, DNA amplification, as a single unit^33^, might be essential to the mass production of bracoviral particles that are injected into parasitized hosts. This could explain the counter selection on the dispersion or loss of these clustered nudivirus genes since the separation of the *Cotesia* and *Microplitis* lineages. The genomic organization stasis is reminiscent of bacterial symbiont genomes, which underwent major rearrangements specifically during the initial steps of association^44^. In the case of the particularly large macrolocus which comprises 77% of virulence genes in *Cotesia* genome, clustering could facilitate the evolution of new virulence genes copies by duplications^13^, and thereby wasp adaptation against host resistance or to new hosts^10,45^ This organization may also promote the transmission of bracovirus virulence genes as a block of co-evolved genes as shown for supergenes involved in butterfly wing patterns^46^ and ant social behavior^47^.

Remarkably, despite their dispersed locations in the wasp genome, bracovirus genes remain synchronically expressed and under conservative selection, thus enabling the production of infectious virus particles. This striking conservation of the viral machinery highlights the paramount importance of the production of viral particles allowing virulence gene transfer to a host, in the evolutionary history of the wasp. Although high nudiviral gene expression and massive particle production underline the continuing existence of a virus entity, it is now considered as a regular secretion by the wasp. Thus, contrary to a vision of obligatory mutualism as an unstable alliance in which partners can become opponents or cheaters^48,49^, this very ancient association shows conflicts disappear when the benefits of the partners converge.

## Methods

### Sampling

The *C. congregata* laboratory strain was reared on its natural host, the tobacco hornworm, *M. sexta* (Lepidoptera: Sphingidae) fed on artificial diet containing nicotine as previously described^50^. *C. sesamiae* isofemale line came from individuals collected in the field in Kenya (near the city of Kitale) and was maintained on *Sesamia nonagrioides*^51^. *C. flavipes* individuals originated from the strain used for biological control against *Diatraea saccharalis* in Brazil^52^. *C. glomerata*, *C. rubecula* and *C. vestalis* laboratory cultures were established from individuals collected in the vicinity of Wageningen in Netherlands, and reared on *Pieris brassicae*, *Pieris rapae* and *Plutella xylostella* larvae, respectively^53,54^. To reduce the genetic diversity of the samples prior to genome sequencing, a limited number of wasps were pooled; for example, only haploid males from a single virgin female were used for Illumina sequencing of *C. congregata* genome, ten female and male pupae originating from a single parent couple were used to generate *C. glomerata* genome, five male pupae originating from a single *C. rubecula* virgin female for *C. rubecula* genome and 40 adult males and 8 adult females from multiple generations of *C. vestalis* cultured in the Netherlands were used. DNAs were extracted from adult wasps and stored in extraction buffer following two protocols. *C. congregata, C. sesamiae* and *C. flavipes* DNA were extracted using a Qiamp DNA extraction kit (Qiagen) with RNAse treatment following the manufacturer’s instructions and eluted in 200 μl of molecular biology grade water (5PRIME). *C. glomerata*, *C. rubecula* and *C. vestalis* DNA was extracted using phenol-chloroform.

### Genome sequencing and assembly

*Cotesia congregata* genome was sequenced combining two approaches: (i) single-end reads and MatePair libraries of 3Kb, 8Kb and 20Kb fragments on 454 GS FLX Titanium platform (Roche) and (ii) paired-end reads of 320bp fragments with HiSeq2000 platform (Illumina). *C. glomerata*, *C. rubecula* and *C. vestalis* DNA libraries were prepared using insert sizes of 300 and 700 bp. For *C. sesamiae* and *C. flavipes* libraries an insert size of 400 bp was selected. These libraries were sequenced in 100 bp paired-end reads on a HiSeq2000 platform (Illumina) at the French National Sequencing Institute (CEA/Genoscope, France) and at the Sequencing Facility of the University Medical Center (Groningen, Netherlands). Reads were then filtered according to different criteria: low quality bases (Q<20) at the read extremities, bases after the second N found in a read, read shorter than 30 bp and reads matching with phiX (Illumina intern control) were removed using in-house software (http://www.genoscope.cns.fr/fastxtend) based on the FASTX-Toolkit (http://hannonlab.cshl.edu/fastx_toolkit/) as described in (https://www.nature.com/articles/sdata201793).

The *C. congregata* genome was generated by *de novo* assembly of 454 reads using GS De Novo Assembler from the Newbler software package v2.8^55^. The consensus was polished using the Illumina data as described in (https://bmcgenomics.biomedcentral.com/articles/10.1186/1471-2164-9-603). Gaps were filled using GapCloser module from the SOAPdenovo assembler (https://www.ncbi.nlm.nih.gov/pmc/articles/PMC3626529/). The genomes of *C. sesamiae*, *C. flavipes, C. glomerata*, *C. rubecula* and *C. vestalis* were assembled with Velvet v1.2.07^55^ using the following parameters: velveth *k*-mer 91-shortPaired-fastq-separate, velvetg-clean yes and specific adapted values for -exp_cov and -cov_cutoff for each species. The genome size was estimated using *k*-mer distribution of raw reads on all *Cotesia* species with 17-mer size using Jellyfish v1.1.11 ^56^.

### Chromosome scale assembly of *C. congregata* genome

#### HiC library preparation

The individual wasps had their gut removed and were immediately suspended after sampling in 30 mL of 1X tris-EDTA buffer and formaldehyde at 3% concentration, then fixed for one hour. 10 mL of glycine at 2.5 M was added to the mix for quenching during 20 min. A centrifugation recovered the resulting pellet for −80°C storage and awaiting further use. The libraries were then prepared and sequenced (2 × 75 bp, paired-end Illumina NextSeq with the first ten bases acting as tags), as previously described^57^ using the *Dpn*II enzyme.

#### Read processing and Hi-C map generation

The HiC read library was processed and mapped onto *Dpn*II fragments of the reference assembly using HiCbox (available at https://github.com/kozullab/HiCbox) with bowtie2^58^ on the back-end (option --very-sensitive-local, discarding alignments with mapping quality below 30). Fragments were then filtered according to size and coverage distribution, discarding sequences shorter than 50 bp or below one standard deviation away from the mean coverage. Both trimmed contact maps were then recursively sum-pooled four times by groups of three, yielding bins of 3^4^= 81 fragments.

#### Contact-based re-assembly

The genome was re-assembled using an updated version of GRAAL (dubbed instaGRAAL) for large genomes on the aforementioned contact maps for 1,000 cycles, as described in^59^. Briefly, the program modifies the relative positions and/or orientations of sequences according to expected contacts given by a polymer model. These modifications take the form of a fixed set of operations (swapping, flipping, inserting, merging, etc.) on the 81-fragment bins. The likelihood of the model is maximized by sampling the parameters with an MCMC method. After a number of iterations, the contact distribution converges and the global genome structure ceases to evolve, at which point the genome is considered reassembled. The process yielded eleven main scaffolds comprising >99% of the bin sequences.

#### Polishing and post-assembly processing

Each scaffold was independently polished by reconstructing the initial contig structure whenever relocations or inversions were found. In addition, previously filtered sequences were reintegrated next to their original neighbors in their initial contig when applicable. The implementation is part of instaGRAAL polishing and available at https://github.com/koszullab/instaGRAAL (run using the –polishing mode).

#### Assembly validation

We performed the validation with QUAST-LG^60^, an updated version of the QUAST analyzer tailored for large genomes. The initial assembly from Illumina short reads was used as reference. The assessed metrics include genomic feature completeness, Nx and NGx statistics as well as global and local misassemblies. In addition, each assembly was assessed for ortholog completeness with BUSCO v3^61^. The reference dataset used for expected gene content was pulled from the OrthoDB (version 9) database for Hymenoptera, comprising 4,415 orthologs.

### Genome annotations

#### Transposable element annotation

Genome annotation was first done on the *C. congregata* reference genome and then exported on the genomes of the five other *Cotesia* species. First, the transposable element annotation was realized following the REPET pipeline comprising a *de novo* prediction (TEdenovo) and an annotation using TE libraries (TEannot)^62^. This annotation was exported into GFF3 files used as mask for the gene annotation.

#### Automated gene annotation

The automated gene prediction and annotation of *C. congregata* genome was done using Gmove (https://github.com/institut-de-genomique/Gmove) integrating different features based on (i) the mapping of Hymenoptera proteins from all hymenopteran genomes available on NCBI (Supplementary Table 6) and UniProt Hymenoptera, (ii) the mapping of RNA-Seq data from *C. congregata*, *C. glomerata*, *C. vestalis* and *C. rubecula*^63,64^, and (iii) *ab initio* genes predictions using SNAP ^65^. The automated annotation of the five other *Cotesia* species was performed using MAKER ^66^ using the same features as for the annotation of *C. congregata* but also including the output automated annotation of *C. congregata*.

#### Automated gene functional annotation

The functional annotation was performed using blastp from BLAST+ v2.5.0^67^ to compare the *C. congregata* proteins to the NCBI non-redundant database (from the 29/06/2014). The ten best hits below an e-value of 1e-08 without complexity masking were conserved. Interproscan v5.13-52.0 ^68^ was used to analyze protein sequences seeking for known protein domains in the different databases available in Interproscan. Finally, we used Blast2GO^69^ to associate a protein with a GO group.

#### Specialist gene annotation

The automated annotations were followed by manual curations, corrections and annotations realized by specialists of each gene family of interest through Apollo^70^. The genomes were available to this consortium through the web portal: https://bipaa.genouest.org/is/parwaspdb/.

#### Genome completeness evaluation

The completeness of the genomes and annotations were evaluated using Benchmarking Universal Single-Copy Orthologs BUSCO v3^61^ using the insecta_odb9 database composed of 1658 genes.

Contigs were searched for similarities against the nonredundant NCBI nucleotide (nt) (release November 2019) and the Uniref90 protein (release November 2019) databases using respectively blastn v2.7.1+^67^ and diamond v0.9.29.130^71^. For both tasks, e-value cutoff was set to 10−25. Taxa were inferred according to the highest-scoring matches sum across all hits for each taxonomic rank in the two databases. Sequencing coverage was deduced after mapping Illumina paired reads to the assembly with Bowtie2 v2.3.4.2^58^. Contigs were then displayed with Blobtools v1.1.1^72^ using taxon-annotated-GC-coverage plots.

### Orthologous genes identification, alignment and phylogeny

Orthologous genes between all genes annotated in the six *Cotesia* species and the four outgroups (*Microplitis demolitor*, *Nasonia vitripennis*, *Apis mellifera* and *Drosophila melanogaster*) were identified using OrthoFinder v1.14^73^. Universal single-copy ortholog genes from BUSCO v3^61^ were extracted for the six *Cotesia* species and the four outgroups, aligned using MAFFT v7.017^74^ and concatenated. The species phylogeny was performed on this alignment composed of 1,058 orthologous for a length of 611 kb using PhyML program^75^ with the HKY85 substitution model, previously determined by jModelTest v2.1^76^ and the branch support were measured after 1,000 bootstrap iterations. The cluster of orthologous genes was used to determine the phylogenetic level of each gene represented in Fig. 2. as follows: genes shared by all species are called shared by all; genes shared by at least nine species among the ten studied species without phylogenetic logical are named “shared by some”; genes shared by only Hymenoptera species and without orthologous gene in *D. melanogaster* are considered as “Hymenoptera specific”; genes shared only by Microgastrinae are named “Microgastrinae specific”; genes shared only by *Cotesia* species and without orthologous genes in any of the outgroup are considered as “Cotesia specific”.

### Synteny analyses

The different synteny analyses were performed on the orthologous genes identified by OrthoFinder v1.14^73^ and by reciprocal blastp from BLAST+ v2.2.28^67^ on the annotated proteins (e-value below 10e^−20^). The correspondence between the genes, the localizations on the scaffold and the figures were realized thanks to a custom R script (R Core Team 2013).

### Evolution of gene families

For OR, P450 and odv-e66 genes manually annotated genes from the reference genome of *C. congregata* were used along with orthologous genes from the five other *Cotesia* species, *M. demolitor*^77^, *N. vitripennis*^78^, *A. mellifera*^79^ to create a phylogeny of each family among Hymenoptera. Protein sequences were first aligned with MAFFT v7.017^74^ and the maximum-likelihood phylogeny was performed with PhyML^75^ using the JTT+G+F substitution model for OR and using HKY80 substitution model for P450 and odv-e66 genes. The branch support was assess using aLRTfor OR and 1,000 bootstraps for P450 and odv-e66 genes.. The trees were then rooted to Orco (OR-coreceptor) clade for OR and t the midpoint for the other. The gene gains and losses along the phylogeny for the different gene families of interest were identified with NOTUNG v2.9 ^80,81^.

### Evolution of single copy genes

To determine evolutionary rates within *Cotesia* genus, single-copy orthologous gene clusters (BUSCO, nudiviral and virulence genes) were first aligned using MACSE^82^ to produce reliable codon-based alignments. From these codon alignments, pairwise dN/dS values were estimated between *C. congregata* and *C. sesamiae*, the two most diverging species in the *Cotesia* phylogeny, with PAML v4.8 using the YN00 program^83^. dN/dS of the different gene categories of interest were then compared using a Kruskal–Wallis test, and Nemenyi-Tests for multiple comparisons were realized with the R package. For the nudiviral genes the dN/dS values were calculated using genes from the 6 species. Orthologous genes from the six *Cotesia* species were aligned as described before and codeml (implemented in PAML v4.8) was used to estimate the M0 dN/dS (free ratio model). This model was compared to a neutral model for which the dN/dS is fixed to 1.

### RNA-Seq analyses

#### Sample preparation, extraction and sequencing

The ovaries and venom glands were extract from females at five pupal stages, i.e. days 2, days 3, days 4, days 5 and at emergence, corresponding to the number of days after the creation of the cocoon and identified following body melanization^84^. Ovaries were pooled by groups of 20 pairs and venom glands by 100 and in duplicates for each condition. Samples were stored in buffer provided in the extraction kit by adding 2-mercaptoethanol to reduce RNA degradation. Extractions were performed using QIAGEN RNeasy kit following manufacturer’s recommendations. Library preparation and sequencing were performed at the Genescope using a paired-end strategy with HiSeq2000 platform (Illumina).

#### Analyses

The pair-end reads from *C. congregata* ovary and venom gland libraries were mapped on the reference genome using TopHat2^85^ with default parameters. Then, featureCounts program from the Subread package^86^ was used to determine fragment counts per genes using default parameters.

To analyze gene expression, the raw fragment counts of ovaries and venom glands samples were first converted to counts per million (CPM) using the edgeR^87^ R-implemented package (R-Core Team 2017). Expressed genes were filtered based on a CPM > 0.4 (corresponding to raw count of 15) in at least two of the libraries incorporated in the analysis (Supplementary Fig. 6 A) and subsequent normalization was performed on CPMs using the edgeR TMM method for Normalization Factor calculation^88^ (Supplementary Fig. 6 B). The reproducibility of replicates was then assessed by Spearman correlation of gene expression profiles based on filtered and normalized CPMs (Supplementary Fig. 6 C).

To examine differential expression between ovary stages and with venom glands a quasi-likelihood negative binomial generalized log-linear model was fitted to the data after estimation of the common dispersion using edgeR. Then, empirical Bayes quasi-likelihood F-tests were performed to identify differentially expressed (DE) genes under chosen contrasts^89^. Finally, F-test p-values were adjusted using False Discovery Rate (FDR) method^90^. Genes were considered as DE whether FDR < 0.05 and fold-change (FC) of expressions between compared conditions was higher or equal to 1.5. Four contrasts were designed between the five successive ovary stages and a control contrast was tested between ovaries and venom glands at wasp emergence stage.

### Data availability

The datasets generated during the current study are available from the National Center for Biotechnology Information (NCBI) at the following BioProject ID: PRJEB36310 for genome raw reads; PRJNA594477 for transcriptome raw reads. Genome database (genomes and annotated genes) are available on the web site BIPAA (Bioinformatic Platform for Agrosystem Arthropods) https://bipaa.genouest.org/is/parwaspdb/. Custom scripts are available from https://github.com/JeremyLGauthier/Script_Gauthier_et_al._2020_NEE.

## Acknowledgements

We thank Paul André Catalayud for providing the pictures of *C. sesamiae* and *C. flavipes* Germain Chevignon for the pictures of *C. congregata* nymphal stages and Juline Herbinière for TEM images of *Cotesia congregata* ovaries. We thank the ADALEP (Adaptation of Lepidoptera) network for the involvement of its members and access to bioinformatic facilities for genome annotation.

## Author information

## Contributions

**DNA/RNA preparation**: J. Gauthier, C. Capdevielle-Dulac, K. Labadie, H. Boulain; F. Consoli, B. L. Merlin;**Sequencing assembly and automatic annotation**: B. Noël, J.-M. Aury, V. Barbe, J. Gauthier, A. Bretaudeau; F. Legeai; ***C. glomerata*, *C. rubecula, C. vestalis* sequencing and first assembly**: J. Van Vugt, H. Smid, J. de Boer, S.Warris, L. Vet; **HiC approach**: L. Baudry, M. Marbouty, R. Koszul; **TEs annotation**: A. Hua-Van (group leader), J. Amselem, I. Luyten; **Bracovirus annotation**: A. Bézier (group leader), J. Gauthier, P. Gayral, K. Musset, T. Josse, D. Bigot, C. Bressac, S. Moreau, E.A. Herniou; **Bracovirus gene evolution and synteny analyses**: J. Gauthier; H. Boulain; **Immune genes annotation**: E. Huguet (group leader), H. Boulain; G. Dubreuil; B. Duvic, J.-M. Escoubas, N. Kremer; **Conserved bracovirus regulatory sequences**: G. Periquet; **Chemosensory genes annotation**: E. Jacquin-Joly (group leader), M. Harry, E. Persyn, N. Montagné, I. Boulogne, M. Sabety, M. Maibeche, T. Chertemps; **Detoxification genes annotation**: G. le Goff (group leader), F. Hilliou, D. Saussiat; **Web annotation online platform**: A. Bretaudeau; F. Legeai; **RNA-Seq analyses**: H. Boulain, J. Gauthier; **Wasp phylogenetic and biological background**: J. B. Whitfield; **Project writing for funding**: S. Dupas, C. L. Kaiser-Arnault, E. Herniou, J.-M. Drezen, H. Smid, L. Vet; **Project coordination**: J. Gauthier and J.-M. Drezen; **Manuscript writing**: J. Gauthier, J.-M. Drezen with contributions from all authors.

## Funding

*C. congregata*, *C. sesamiae*, *C. flavipes* genomes sequencing were funded by French National Research Agency ANR (ABC Papogen project ANR-12-ADAP-0001 to L. Kaiser-Arnaud). *C. rubecula*, *C. glomerata*, *C. vestalis* genomes sequencing were funded by NWO EcoGenomics grant 844.10.002 to L.E.M. Vet, NWO VENI grant 863.07.010 and Enabling Technology Platform Hotel grant to L.E.M. Vet. HiC approach was funded by ERC project 260822 to R. Koszul. *C. congregata* transcriptomic analysis was funded by APEGE project (CNRS-INEE) to J.-M. Drezen. J. Gauthier thesis was funded by ANR and Region Centre-Val de Loire. Collaboration between French and Netherland laboratories was funded by French ministry of foreign affairs and “Nuffic” (“VanGogh” Project to J-M Drezen and L.E.M. Vet).

## Ethics declarations

### Competing interests

The authors declare no competing interests.

## Additional information

Publisher’s note: Springer Nature remains neutral with regard to jurisdictional claims in published maps and institutional affiliations.

## Supplementary information

Cotesia_genomes_Supplementary_data.pdf

