## Supplementary data for "Chromosomal resolution reveals symbiotic virus colonization of parasitic wasp genomes"

### Chromosomal resolution of a parasitic wasp genome reveals the colonisation of its symbiotic virus

SUPPLEMENTARY INFORMATION (NOTES, TABLES, FIGURES)

A. Global analyses

1. Genome sequencing and assembly
2. Chromosome scale assembly of *C. congregata* genome
3. Genome annotations
4. RNAseq analyses

B. Phylogenetic considerations

C. Bracovirus

D. Immunity

E. Chemoreceptor

F. Detoxification

References

#### A. Global analyses

##### 1. Genome sequencing and assembly

**Supplementary Table 1.** Sequencing and assembly statistics.

| Statistics | *C. congregata* | | | *C. rubecula* | *C. glomerata* | | *C. vestalis* | *C. flavipes* | *C. sesamiae* |
|---|---|---|---|---|---|---|---|---|---|
| Sequencing technology | 454 single end | 454 mate-pair (3,8, 20kb) | Illumina | Illumina | Illumina | PacBio | Illumina | Illumina | Illumina |
| Coverage | 20 X | 5 X | 132 X | 237 X | 196 X | 4 X | 216 X | 246 X | 242 X |
| # Scaffolds | 3,140 | | | 35,383 | 50,739 | | 31,915 | 18 696 | 13,504 |
| N50 scaffold | 1,122.4 kb | | | 12.8 kb | 9.1 kb | | 14.6 kb | 20.3 kb | 26.6 kb |
| L50 scaffold count | 48 | | | 4,490 | 6,813 | | 2,549 | 2,071 | 1,699 |
| Mean scaffold size | 65,892 | | | 6,120 | 4,800 | | 5,521 | 8,28 | 8,952 |
| Median scaffold size | 3,331 | | | 2,940 | 2,497 | | 2,047 | 3,278 | 2,697 |
| Longest scaffold | 4,757 kb | | | 156 kb | 197 kb | | 561 kb | 166 kb | 228 kb |
| Total length | 206.9 Mb | | | 216.5 Mb | 243.5 Mb | | 176.2 Mb | 154.8 Mb | 165.9 Mb |
| %N | 6.68 | | | 3.97 | 4.62 | | 3.75 | 1.78 | 2.74 |
| GC content (%) | 28.21 | | | 29.10 | 28.73 | | 29.26 | 29.62 | 29.35 |

**Supplementary Table 2.** Statistics used for the genome size estimation (M: mean K-mer coverage, N: genome coverage, K: K-mer size, L: mean read length, N=(M*L)/(L-K+1)) and genome size measured by cytometry.

| | | *C. congregata* | *C. rubecula* | *C. glomerata* | *C. vestalis* | *C. flavipes* | *C. sesamiae* |
|---|---|---|---|---|---|---|---|
| k-mer counting estimation | M | 149 | 160 | 106 | 156 | 176 | 171 |
| | K | 17 | 17 | 17 | 17 | 17 | 17 |
| | L | 98.94 | 91.07 | 79.32 | 91.55 | 98.84 | 98.77 |
| | N | 177.74 | 97.05 | 66.39 | 94.51 | 209.99 | 204.05 |
| | Genome size estimation | 215.9 Mb | 204.7 Mb | 160.8 Mb | 129.8 Mb | 178.1 Mb | 193.2 Mb |
| | Genome assembled size | 206.9 Mb | 216.5 Mb | 243.5 Mb | 176.2 Mb | 154.8 Mb | 165.9 Mb |
| | Cytometry measure | n.a. | 220 Mb | 298 Mb | 189 Mb | n.a. | n.a. |

#### 2. Chromosome scale assembly of *C. congregata* genome

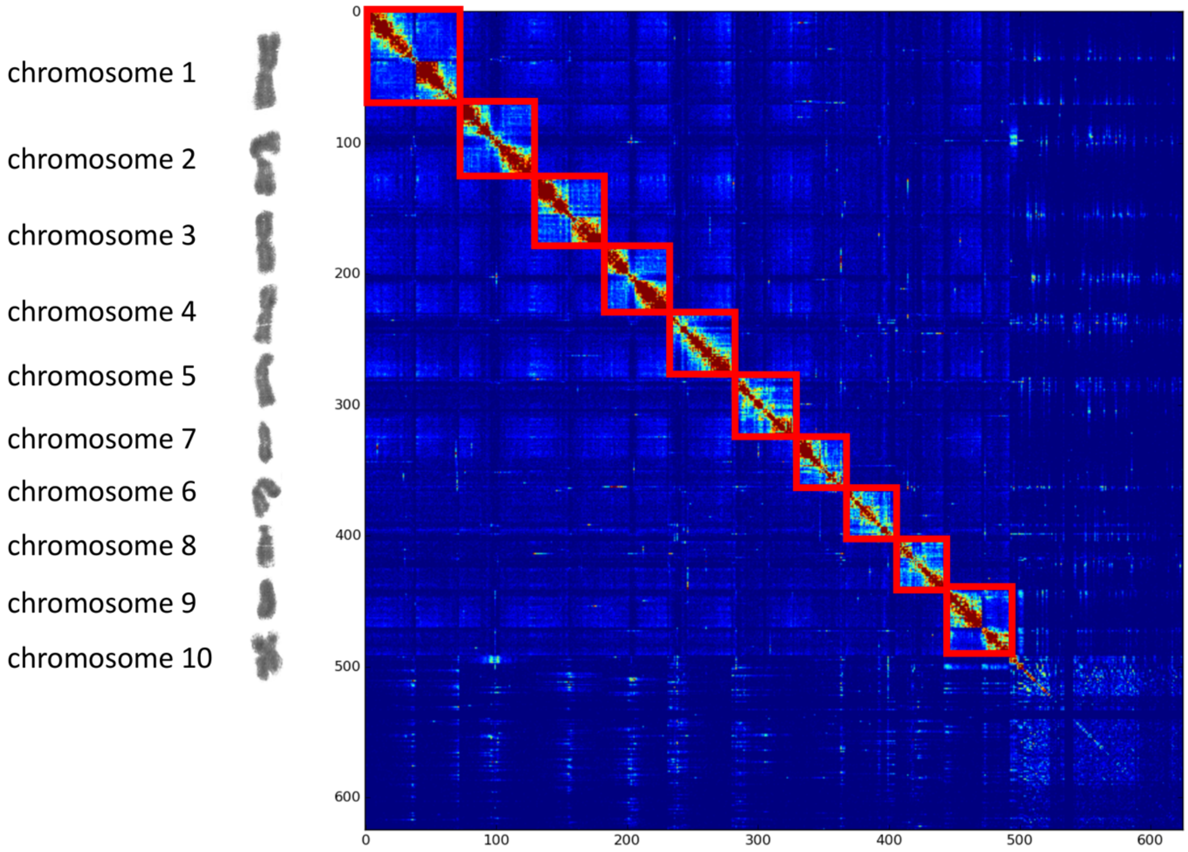

**Supplementary Figure 1**. Contact matrix between each fragment used for the GRAAL assembly, from blue the most distant to red the closest fragments. Along the two axis the fragments are organized according to this distance. The red frames have been added to visualize the chromosomes separations. The assignation of each chromosome has been performed using the chromosome length and centromere positions but also including sequences used for in situ hybridization performed by Belle and colleagues[1] (the chromosome pictures have been extracted from karyotype performed in this study).

**Supplementary Table 3**. Comparative statistics of *C. congregata* assembly strategies.

| Statistics | *C. congregata* scaffolds | *C. congregata* chromosomes |
|---|---|---|
| # Scaffolds | 3,140 | 1,790 |
| N50 scaffold | 1,122.4 kb | 20,028.0 kb |
| L50 scaffold count | 48 | 5 |
| Mean scaffold size | 65,892 | 111,241 |
| Median scaffold size | 3,331 | 1,708 |
| Longest scaffold | 4,757 kb | 29,601 kb |
| Total length | 206.9 Mb | 199.1 Mb |
| %N | 6.68 | 6.71 |
| GC content (%) | 28.21 | 27.97 |

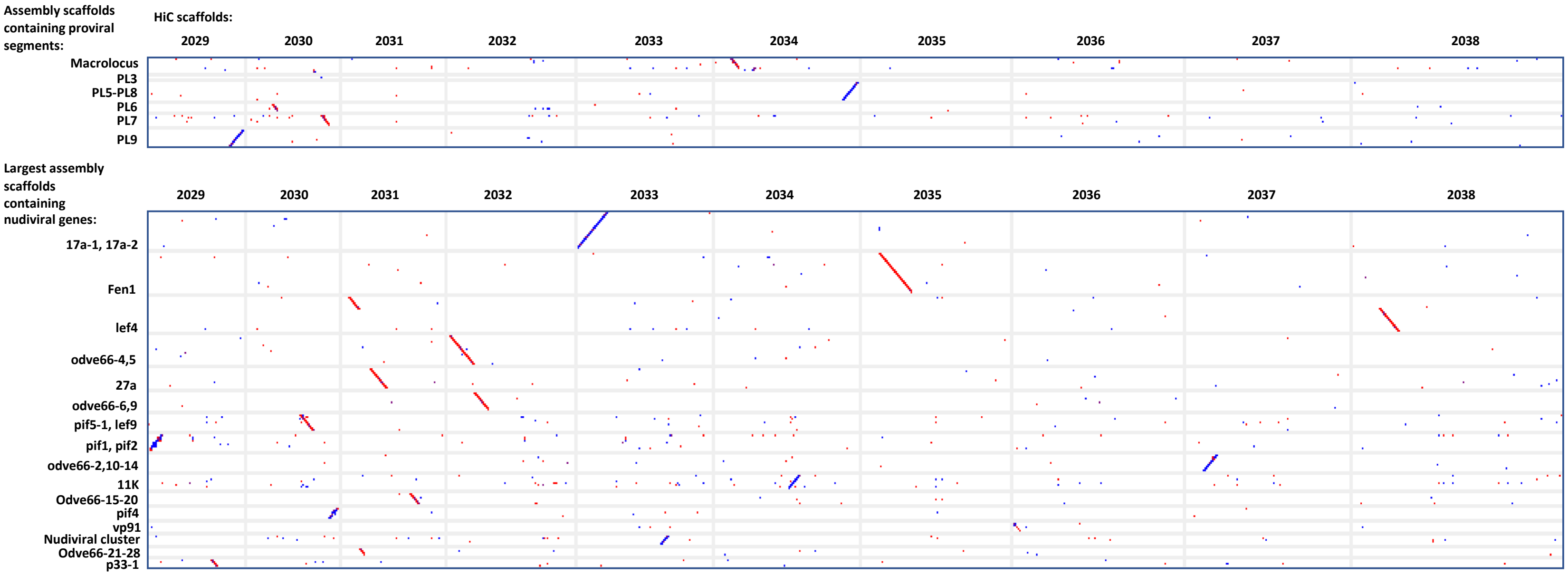

**Supplementary Figure 2.** Dot plot: assembly scaffolds/HiC scaffolds. Assembly scaffolds are confirmed by HiC scaffolds (except for the *lef4*-containing scaffold which is split in two pieces in HiC).

**Supplementary Table 4.** Chromosome size and centromere positions.

| Chromosome | Chromosome size (bp) | Centromere start interval | Centromere stop interval | Centromere size [min - max ] |
|---|---|---|---|---|
| C1 | 29,601,432 | [ 12,811,254 - 13,152,523 ] | [ 13,793,916 - 14,111,909 ] | [ 641,393 - 1,300,655 ] |
| C2 | 23,655,861 | [ 8,494,020 - 8,801,912 ] | [ 13,183,609 - 13,558,132 ] | [ 4,381,697 - 5,064,112 ] |
| C3 | 22,997,154 | [ 8,843,013 - 9,160,875 ] | [ 12352173 - 12654164 ] | [ 3,191,298 - 3,811,151 ] |
| C4 | 20,820,356 | [ 8,962,848 - 9,283,572 ] | [ 12,33,9695 - 12,649,662 ] | [ 3,056,123 - 3,686,814 ] |
| C5 | 20,027,981 | [ 5,848,332 - 6,120,763 ] | [ 9,521,997 - 9,837,552 ] | [ 3,401,234 - 3,989,220 ] |
| C7 | 18,774,300 | [ 13,936,098 - 14,262,658 ] | [ 16,606,145 - 16,890,888 ] | [ 2,343,487 - 2,954,790 ] |
| C6 | 17,819,366 | [ 7,190,494 - 7,542,794 ] | [ 8,206,837 - 8,525,856 ] | [ 664,043 - 1,335,362 ] |
| C8 | 14,366,765 | [ 8,317,315 - 8,636,383 ] | [ 9,295,605 - 9,620,272 ] | [ 659,222 - 1,302,957 ] |
| C9 | 13,378,970 | [ 1,530,552 - 1,939,377 ] | [ 3,558,308 - 3,831,838 ] | [ 1,618,931 - 2,301,286 ] |
| C10 | 12,702,143 | [ 3,965,924 - 4,320,652 ] | [ 6,170,807 - 6,462,798 ] | [ 1,850,155 - 2,496,874 ] |

#### 3. Genome annotations

##### Transposable elements

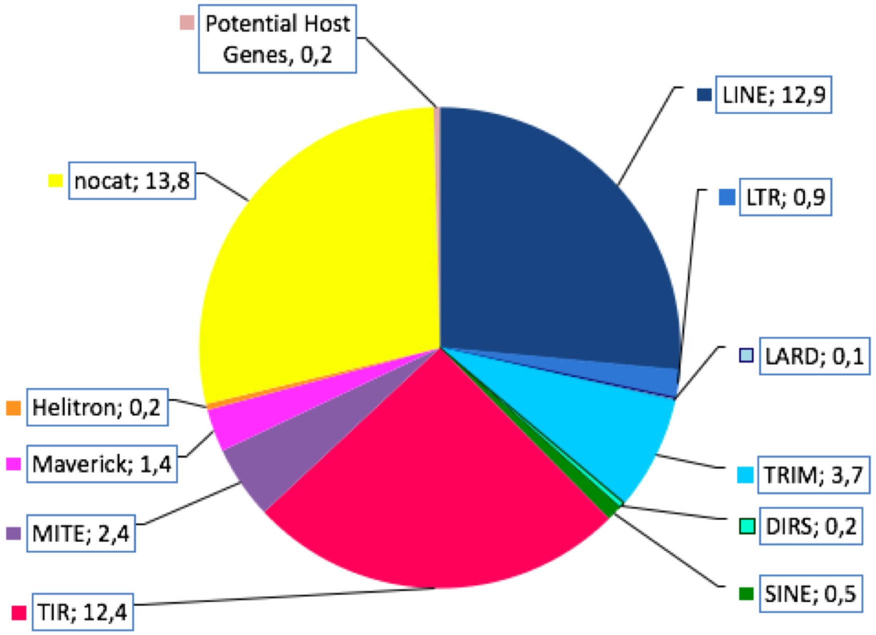

**Supplementary Figure 3.** Pie plot of transposable elements organized by families.

**BV genes potentially originating from transposable elements**

*Analysis of the BV26 related sequences*
The annotation of transposable elements by REPET revealed that the BV26 corresponded to a MITE (Miniature Inverted-repeat Transposable Element), 1606 bp long with 616 nt TIR (Terminal Inverted Repeats). A preliminary search detected 13 very similar copies of this sequence in the genome of *C. congregata*. The BV26 ORF presented no homology with TE proteins (blastx search against the RepBase protein database (RepBase20.05_REPET edition, https://www.girinst.org/) with default parameters.
*Detection of longer related elements*
We then used the 60 first nt of the TIR of this sequence in a blastn search against the genome of *C. congregata* and were able to detect and extract 197 sequences bordered by convergent TIRs (home-made script). Those sequences were blasted against the repbase protein database (RepBase20.05_REPET edition) (blastx, -evalue 1e-4). 49 sequences gave hits with TE proteins, among which 24 (46 different hits) had homology with Sola2 transposases (Class II elements[2]).
The presence of TIRs was rechecked on the sequences, with IRF[3]. We kept 146 sequences containing no N in the sequence, and the sequences were clustered using usearch v5.1 ([4] -id=0.8). Among the 31 clusters found, 6 of them, containing most of the sequences presenting homology with Sola2 elements and characterized by TIRs similar to the TIRs observed in BV26. Two other clusters corresponded to MITE sequences with no protein homology, but long TIRs, and containing amplified copies. This suggested that the MITE sequences could have derived of Sola2 elements after internal deletions. These 8 clusters corresponded to 82 sequences and were kept for further analyses (Supplementary Figure 4 A). Other clusters, presenting either terminal sequences not corresponding to the BV26 TIRs (internal parts of TIRs of another TE family), or sequences of various sizes that may correspond to relics having inserted foreign sequences, were excluded from the analyses. The presence of a large number of sequences varying in size, suggest that this element is ancient.
*Characterization of insertions sites*
In order to better characterize this element, the flanking sequences of the selected copies (200 nt each side) were used in a blastn search against the *C. congregata* genomes to detect paralogous (repetitive) sequences without Sola2 insertions. Comparison of these paralogous sequences with Sola2 corresponding insertion sites allowed us to propose that the TIRs of the elements start with 4 Gs, and that the element inserts into TA-rich sequences, creating a 4-bp target site duplication as expected for Sola2 elements[2]. For a number of cases, insertion sites occurred in very TA-rich microsatellites-like TA sequences and the paralogous sites displayed a variability in the number of TA repeats, making the TSD difficult to identify with confidence. Some examples of clear-cut cases are shown in Supplementary Figure 4 B.
*Characterization of Sola2 sequences*
The 46 sequence fragments presenting homology to Sola2 transposases were translated (and merged when corresponding to different fragment of the same copy) and aligned with selected sequences of Sola2 from RepBase, plus 3 sequences of the related Sola3 element used as outgroup (MAFFT v7[5]) The alignment was truncated to keep only the 250 AA-long best conserved part (AA 240 to 520 of Sola2-1_Nvi) for a phylogenetic analysis using FastTree[6] 2 sequences were removed (deleted over this conserved region). The resulting tree suggests a monophyletic origin of the *C. congregata* Sola2 sequences. Our knowledge of Sola2 elements remains fragmentary, only few genomes having been screened. Closest sequences in Repbase are found in the related parasitoid

*Nasonia vitripennis,* and two ant species (Supplementary Figure 4 C). Potentially active sequences (long ORFs, not stop codons or frameshift, with one intron) could be identified in clusters 6 and 16.

*Distribution among other genomes*

Finally, we searched for related sequences in 5 other *Cotesia* species and in other genomic sequences from Hymenoptera and Lepidoptera. For this, coding fragments from one sequence per cluster were blasted (blastn, -evalue 1e-10) against the genome assemblies of these two insect orders, available in NCBI (by October 14$^{th}$, 2019), representing 309 assemblies of Hymenoptera (278 species) and 204 assemblies of Lepidoptera (64 species). Hits representing the same sequence were merged using a home-made script (maximum distance for merging 2 hits: 2000 nt, and minimum final size: 100 nt). The distribution of the elements among the different species tested, as well as the distribution of the hits among the different assemblies for each superfamily of organisms is shown on Supplementary Figure 4. Phylogenetic relationships between taxa are derived from[7,8]. Among Hymenoptera, similar sequences were detected in 71 species, mostly within the Parasitoida superfamily, with some species displaying a high number of sequences (in Chalcidoidea and Ichneumonoidea), as well as in some other groups, especially in ants (Formicoidea). 17 Lepidoptera species also exhibited some related sequences, but always in relatively low copy numbers. We computed the percentage of identity given by the blastn search, between those hits and the *C. congregata* query sequences (Supplementary Figure 4 E). The most abundant cluster in *Cotesia* was Cluster 7 (See Supplementary Table 5). For this cluster, the average percentages of identity were higher for *Cotesia* genomes than for other Hymenoptera or for Lepidoptera. However, sequences in Lepidoptera had a higher identity than Hymenoptera sequences. The best Lepidoptera hits (>90% over 300 nt) were found in *Papillio machaon*. This may reflect some past events of horizontal transfers between these two orders of insects[9]. Other clusters presented contrasted patterns suggesting absence of close sequences in some *Cotesia* genomes, and/or existence of other Sola2 cluster not found in *C. congregata*.

*Distribution of other repeated sequences associated to BV genes*

The same blastn procedure was applied to all the consensus sequences corresponding to repetitive sequences. The number of hits is shown for each consensus on Supplementary Table 5.

**Supplementary Table 5.** Distribution of reconstituted hits for different repeated sequences related to expressed BV genes.

| | Sola2 ORF (BV26) | | | | | | MITE BV26 | | REPET consensus sequences | | | | | | | | |
|---|---|---|---|---|---|---|---|---|---|---|---|---|---|---|---|---|---|
| | Cluster 6 | Cluster 7 | Cluster 8 | Cluster 15 | Cluster 16 | Cluster 20 | Cluster 18 | Cluster 23 | BV14.3 | BV2.3 | BV21.1 | BV26 | BV3.5 | BV3.7 | BV30.5B | BV4.4 | BV7 |
| *Cotesia congregata* | 23 | 70 | 10 | 3 | 11 | 13 | 231 | 548 | 256 | 37 | 100 | 717 | 224 | 108 | 53 | 313 | 889 |
| *Cotesia rubecola* | 17 | 130 | 64 | 12 | 9 | 9 | 297 | 342 | 365 | 24 | 80 | 516 | 145 | 102 | 117 | 479 | 539 |
| *Cotesia glomerata* | 10 | 112 | 36 | 15 | 4 | 9 | 192 | 396 | 238 | 30 | 69 | 502 | 225 | 117 | 148 | 469 | 533 |
| *Cotesia vestalis* | 20 | 74 | 26 | 15 | 8 | 9 | 106 | 255 | 137 | 5 | 32 | 345 | 87 | 32 | 52 | 161 | 419 |
| *Cotesia sesamiae* | 23 | 77 | 30 | 4 | 7 | 4 | 109 | 424 | 122 | 3 | 39 | 486 | 106 | 20 | 68 | 213 | 479 |
| *Cotesia flavipes* | 25 | 79 | 43 | 9 | 6 | 5 | 142 | 369 | 98 | 2 | 36 | 476 | 123 | 20 | 43 | 251 | 480 |
| Hymenoptera (w/o *Cotesia*) | 523 | 1074 | 247 | 747 | 453 | 171 | 156 | 674 | 332 | 19 | 65 | 190 | 190 | 81 | | 325 | |
| Lepidoptera | 2 | 9 | 1 | 10 | 15 | 1 | 32 | 147 | | | | | | | | | |
| | 643 | 1625 | 457 | 815 | 513 | 221 | 1265 | 3155 | 1548 | 120 | 421 | 3232 | 1100 | 480 | 481 | 2211 | 3339 |

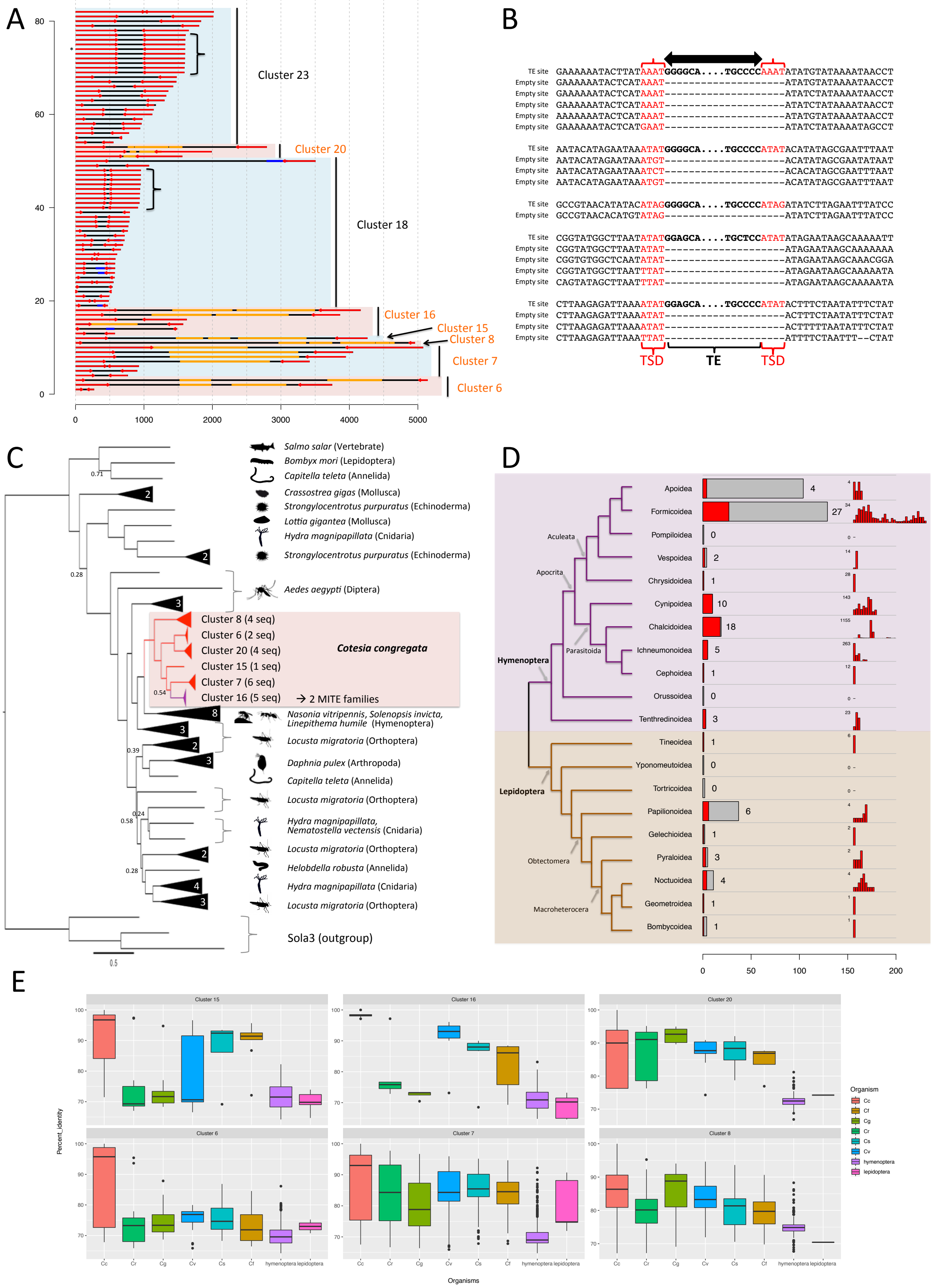

A
80
60
40
20
0
0
1000
2000
3000
4000
5000
Cluster 23
Cluster 20
Cluster 18
Cluster 16
Cluster 15
Cluster 8
Cluster 7
Cluster 6
B
TE site GAAAAAATACTTATAAATGGGGCA....TGCCCCAAATATATGTATAAAATAACCT
Empty site GAAAAAATACTCATAAAT--------------------ATATGTATAAAATAACCT
Empty site GAAAAAATACTCATAAAT--------------------ATATCTATAAAATAACCT
Empty site GAAAAAATACTCATAAAT--------------------ATATCTATAAAATAACCT
Empty site AAAAAAATACTCATAAAT--------------------ATATGTATAAAATAACCT
Empty site GAAAAAATACTCATGAAT--------------------ATATCTATAAAATAGCCT
TE site AATACATAGAATAAATATGGGGCA....TGCCCCATATACATATAGCGAATTTAAT
Empty site AATACATAGAATAAATGT--------------------ACATATAGCGAATATAAT
Empty site AATACATAGAATAAATCT--------------------ACACATAGCGAATTTAAT
Empty site AATACATAGAATAAATGT--------------------ACATATAGCGAATTTAAT
TE site GCCGTAACATATACATAGGGGGCA....TGCCCCATAGATATCTTAGAATTTATCC
Empty site GCCGTAACACATGTATAG--------------------ATATCTTAGAATTTATCC
TE site CGGTATGGCTTAATATATGGAGCA....TGCTCCATATATAGAATAAGCAAAAATT
Empty site CGGTATGGCTTAATATAT--------------------ATAGAATAAGCAAAAAAA
Empty site CGGTGTGGCTCAATATAT--------------------ATAGAATAAGCAAACGGA
Empty site CGGTATGGCTTAATTTAT--------------------ATAGAATAAGCAAAAATA
Empty site CAGTATAGCTTAATTTAT--------------------ATAGAATAAGCAAAAATA
TE site CTTAAGAGATTAAAATATGGAGCA....TGCCCCATATACTTTCTAATATTTCTAT
Empty site CTTAAGAGATTAAAATAT--------------------ACTTTCTAATATTTCTAT
Empty site CTTAAGAGATTAAAATAT--------------------ACTTTTTAATATTTCTAT
Empty site CTTAAGAGATTAAATTAT--------------------ATTTTCTAATTT--CTAT
TSD
TE
TSD
C
0.71
0.28
0.54
0.39
0.24
0.58
0.28
0.5
Salmo salar (Vertebrate)
Bombyx mori (Lepidoptera)
Capitella teleta (Annelida)
Crassostrea gigas (Mollusca)
Strongylocentrotus purpuratus (Echinoderma)
Lottia gigantea (Mollusca)
Hydra magnipapillata (Cnidaria)
Strongylocentrotus purpuratus (Echinoderma)
Aedes aegypti (Diptera)
Cluster 8 (4 seq)
Cluster 6 (2 seq)
Cluster 20 (4 seq)
Cluster 15 (1 seq)
Cluster 7 (6 seq)
Cluster 16 (5 seq)
→ 2 MITE families
Cotesia congregata
Nasonia vitripennis, Solenopsis invicta, Linepithema humile (Hymenoptera)
Locusta migratoria (Orthoptera)
Daphnia pulex (Arthropoda)
Capitella teleta (Annelida)
Locusta migratoria (Orthoptera)
Hydra magnipapillata, Nematostella vectensis (Cnidaria)
Locusta migratoria (Orthoptera)
Helobdella robusta (Annelida)
Hydra magnipapillata (Cnidaria)
Locusta migratoria (Orthoptera)
Sola3 (outgroup)
D
Hymenoptera
Aculeata
Apocrita
Parasitoida
Lepidoptera
Obtectomera
Macroheterocera
Apoidea 4
Formicoidea 27
Pompiloidea 0
Vespoidea 2
Chrysidoidea 1
Cynipoidea 10
Chalcidoidea 18
Ichneumonoidea 5
Cephoidea 1
Orussoidea 0
Tenthredinoidea 3
Tineoidea 1
Yponomeutoidea 0
Tortricoidea 0
Papilionoidea 6
Gelechioidea 1
Pyraloidea 3
Noctuoidea 4
Geometroidea 1
Bombycoidea 1
0
50
100
150
200
E
Cluster 15
Cluster 16
Cluster 20
Cluster 6
Cluster 7
Cluster 8
Percent_identity
Organisms
Cc
Cr
Cg
Cv
Cs
Cf
hymenoptera
lepidoptera
Organism

**Supplementary Figure 4.** Characteristics of Sola2 elements from *C. congregata.* **A** Structure of the 82 sequences belonging to the 8 main clusters. The x-axis is the length in nt. Six clusters (orange) contained sequences with homology to Sola2 transposase (orange part). Blue parts indicate homology with other TEs from RepBase. The two more abundant correspond to 2 MITE clusters and copies resulting from amplification (same size and sequences) are highlighted by brackets. Red arrows represent the TIRs of the elements. The asterisk indicates the initial copy corresponding to BV26. **B** Graph showing the target site duplication (4-nt, TA rich) deduced from the comparison of some Sola2 insertions sites with paralogous empty sites. As for typical Sola2 element, TIRs start by G-rich sequences. **C** Phylogenetic tree showing the relationships between Sola2 translated sequences from *C. congregata* (framed branches) and other Sola2 transposases from RepBase. Black triangles group TE consensus sequences from the same species (number of sequences is indicated). Open triangles represent consensus sequences of Sola3 elements (RepBase) used as an outgroup. Numbers on the branches are the robustness (local bootstraps) as implemented in FastTree. One group (purple) may be at the origin of the 2 MITEs families (based on TIR similarities). **D** Distribution of related Sola2 elements in genomes of hymenoptera and lepidoptera, available in NCBI. Grey bars represent the number of species for which genome sequences are available, and tested for the presence of Sola2 sequences. The red parts of the bars correspond to the number of species in which blastn hits have been found (numbers of species are indicated on the right of the bars. The number of different hits for each genome assembly is shown at the far right, the numbers beside correspond to the maximum number of hits found in one assembly (the highest bar in each barplot). Note that the number of assemblies can exceed the number of positive species (several genome assemblies for one species). **E** Boxplots showing the distribution of the percentage of identity between the *C. congregata* Sola2 sequences (ORF) (1 copy per cluster) and the blastn hits found in the various genomes. Cc: *C. congregata*, Cr: *C. rubecula*, Cg: *C. glomerata*, Cv: *C. vestalis*, Cs: *C. sesamiae*, Cf: *C. flavipes.*

**Supplementary Table 6.** Databases used during the annotation process.

| | Species | Version | Database | Link |
|---|---|---|---|---|
| Automated annotation | *Apis mellifera* | v3.2 | Beebase | http://hymenopteragenome.org/beebase/ |
| | *Nasonia vitripennis* | v1.2 | NasoniaBase | http://www.hymenopteragenome.org/nasonia/ |
| | *Acromyrmex echinatior* | v.3.8 | AntGenomes | http://antgenomes.org/ |
| | *Atta cephalotes* | v1.0 | AntGenomes | http://antgenomes.org/ |
| | *Camponotus floridanus* | v1.0 | AntGenomes | http://antgenomes.org/ |
| | *Harpegnathos saltator* | v1.0 | AntGenomes | http://antgenomes.org/ |
| | *Linepithema humile* | v1.2 | AntGenomes | http://antgenomes.org/ |
| | *Pogonomyrmex barbatus* | v1.2 | AntGenomes | http://antgenomes.org/ |
| | *Solenopsis invicta* | v2.2.3 | AntGenomes | http://antgenomes.org/ |
| Functional annotation | *Acromyrmex echinatior* | v.3.8 | AntGenomes | http://antgenomes.org/ |
| | *Apis mellifera* | v3.2 | Beebase | http://hymenopteragenome.org/beebase/ |
| | *Drosophila melanogaster* | v6.03 | Flybase | http://flybase.org/ |
| | *Nasonia vitripennis* | v1.2 | NasoniaBase | http://www.hymenopteragenome.org/nasonia/ |
| | *Manduca sexta* | OGS 2.0_20140407 | ManducaBase | http://agripestbase.org/manduca |

**Supplementary Table 7.** Genome annotation statistics

| Statistics | *C. congregata* | *C. rubecula* | *C. glomerata* | *C. vestalis* | *C. flavipes* | *C. sesamiae* |
|---|---|---|---|---|---|---|
| Number of predicted genes | 14,140 | 22,795 | 23,498 | 19,239 | 17,381 | 17,785 |
| Mean gene size (bp) | 5,919.47 | 2,852.67 | 2,669.74 | 2,840.57 | 3,238.76 | 3,466.08 |
| Number of exons | 69,667 | 96,948 | 95,98 | 84,368 | 85,318 | 88,955 |
| Mean number of exon by gene | 4.93 | 4.25 | 4.08 | 4.39 | 4.91 | 5.00 |
| Mean exon size (bp) | 365.33 | 310.30 | 304.81 | 306.69 | 306.74 | 313.94 |
| % coding sequence in genome | 12.30 | 13.89 | 12.01 | 14.69 | 16.90 | 16.83 |
| Number of gene without intron | 1630 | 2815 | 2919 | 2330 | 1803 | 1687 |
| Mean intron size (bp) | 1047.62 | 469.23 | 459.88 | 439.81 | 441.39 | 471.76 |

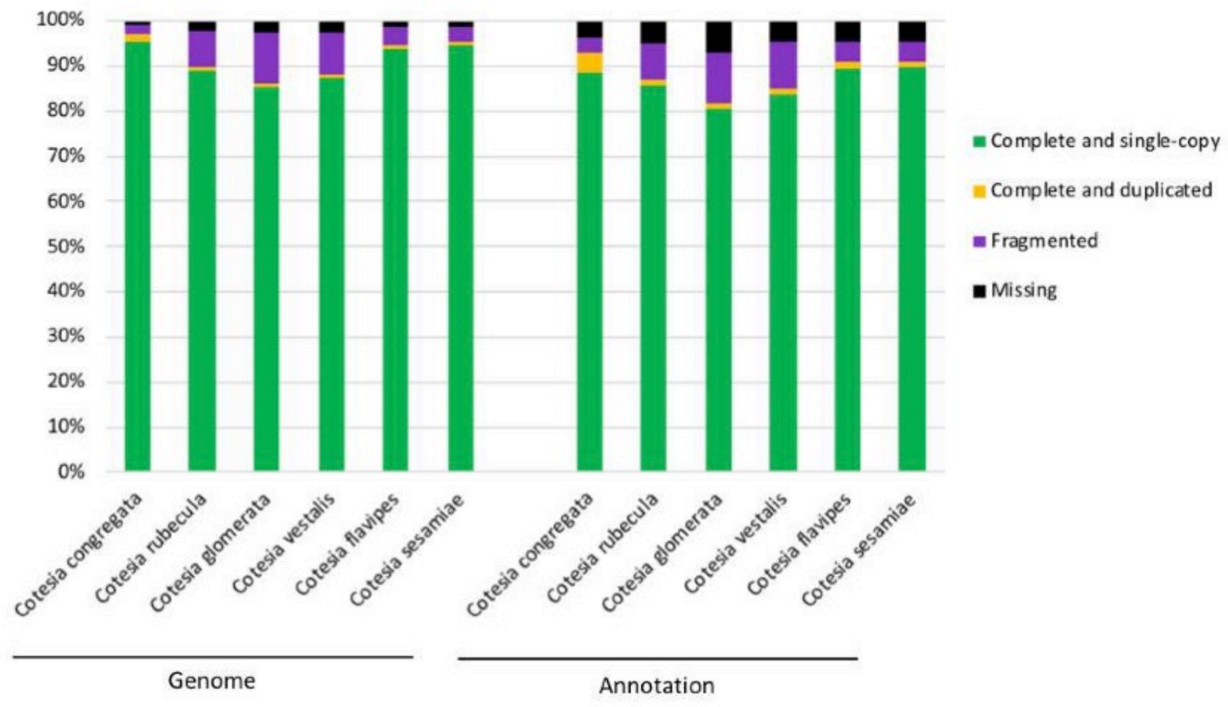

**Supplementary Figure 5.** BUSCO results obtained on genome and annotation of each *Cotesia* species.

#### 4. RNA-Seq analyses

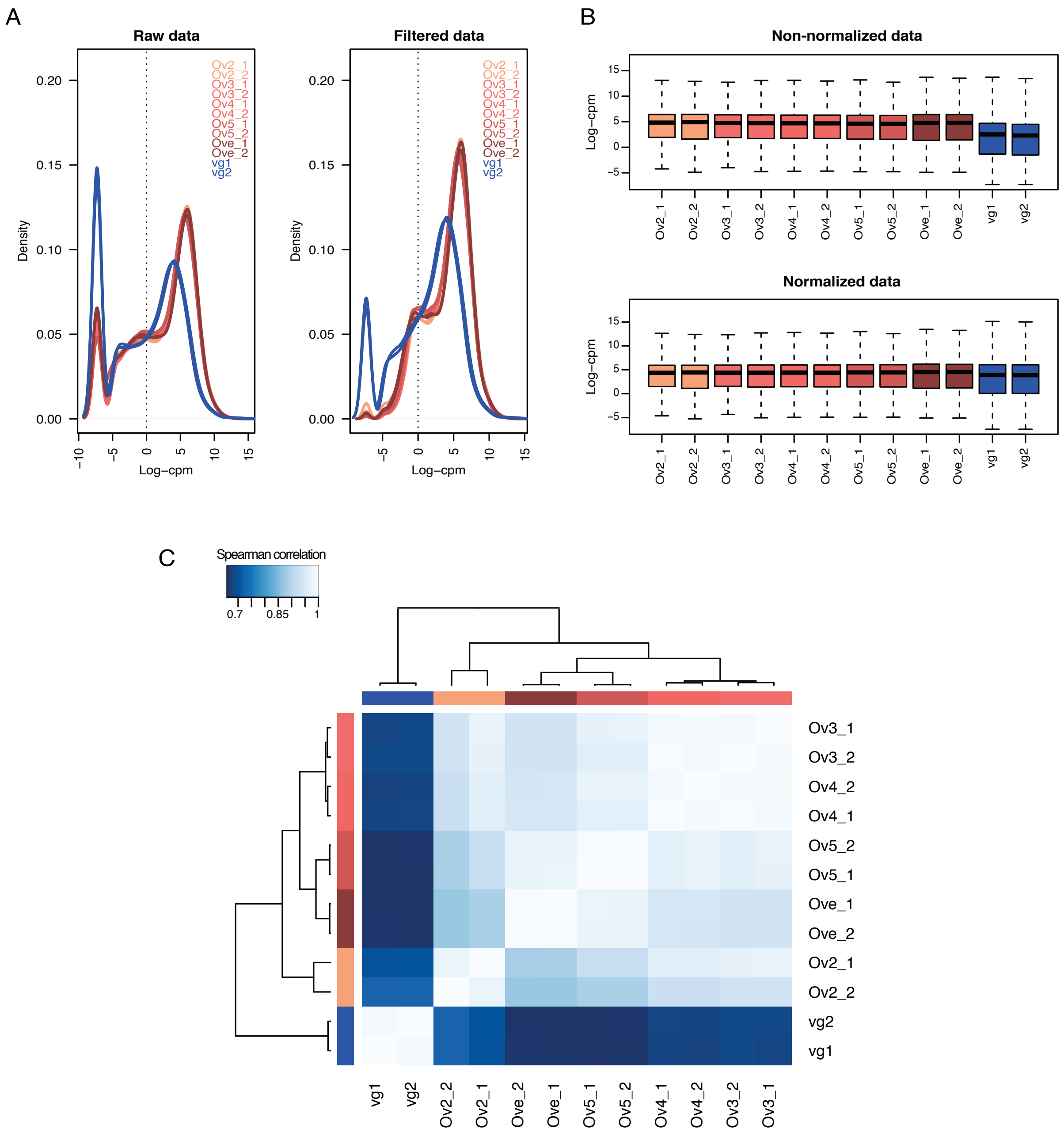

**Supplementary Figure 6.** Gene expression analysis of ovaries and venom glands of *Cotesia congregata.* **A.** Density plots representing expression levels of all expressed genes (Raw data: 13,607 genes) and after filtering the genes that do not show CPM > 0.4 in at least 2 libraries on the 12 analyzed libraries (Filtered data: 11,216 genes). **B.** Expression level distribution of the genes (filtered data) before and after CPM normalization using TMM method in edgeR. **C.** Heatmap of Spearman correlation between the 12 analyzed libraries. The unsupervised clustering did not reveal discrepancies between biological replicates, then all libraries were retained for further analyses. Ov2, Ov3, Ov4 and Ov5 represent ovary samples collected at different larval stages. Ove and vg respectively refer to ovaries and venom glands from adult wasp.

#### B. Phylogenetic considerations

*Monophyly of the bracovirus-bearing lineage*

Bracoviruses have so far been identified from the Microgastrinae (many species), Cardiochilinae (especially *Toxoneuron nigriceps*), Miracinae, Mendesellinae[10] and Cheloninae (several genera, but especially *Chelonus inanitus*). These all belong to what is commonly referred to today as the "microgastroid assemblage". To date, no sample from this group of subfamilies has ever failed to reveal an associate bracovirus (although of course the sampling is still rather sparse). A grouping within the braconid wasps very roughly corresponding to this assemblage, as currently conceived, has been recognized since at least the 1960's, if not before, although its current composition was not well understood (and to some extent is still not fully supported by strong evidence for a few possibly peripheral taxa). A close relationship between Microgastrinae, Cardiochilinae and Miracinae (sometimes also Adeliinae) was clear, but it was not until later studies showed that Adeliinae actually belong within the Cheloninae[11,12] and two new related subfamilies, the South African Khoikhoiinae[13] and the largely Neotropical Mendesellinae[14] were described, that the core microgastroid taxa were identified. Quicke and Van Achterberg[15], in a morphology-based phylogenetic study of Braconidae, found Dirrhopinae and Ichneutinae, both suspected relatives, to be part of the same assemblage, but as its earliest-diverged lineages. To date neither Dirrhopinae nor Ichneutinae (both relatively uncommon groups) have been surveyed for bracoviruses.

Monophyly of the bracovirus-bearing lineage of braconid wasps was first established using morphological and molecular data by[16] and subsequently confirmed by a number of molecular phylogenetic studies (e. g.[17-21]. The (presumably single) origin of the bracoviruses in these wasp groups has been confirmed as from ancestral nudiviruses[10], whose association with insects extends back much earlier[22].

*Relationships among microgastroid subfamilies*

Initially, relationships among the microgastroid subfamilies were established primarily using morphological data[14]. As additional subfamilies in the assemblage were described, and molecular data became available[16,17,20,21], the relationships shown in Figure 1 began to be confirmed. There still remains some uncertainty about the exact relationships among the subfamilies, and about the relationship of Ichneutinae and Dirrhopinae to the core microgastroids, but generally speaking this is one of the best-established groupings within the braconid wasps. One constant is that Cheloninae (including Adeliini) is likely to be sister to the clade including Mendesellinae, Khoikhoiinae, Cardiochilinae, Miracinae and Microgastrinae, diverging from the others roughly 100 million years ago.

*Relationships within Microgastrinae*

The subfamily Microgastrinae is one of the most species-rich parasitoid groups on earth, with an estimated fauna of 17-46,000 species worldwide, based on various field-study extrapolations from the described species. Genus level systematics of the group remains in flux – currently 81 genera are recognized[23], several of them containing more than 1,000 species. As a result of this diversity, and also due to the genera having apparently evolved in a rapid burst roughly 50 million years ago[19,20,24,25], the higher-level phylogeny within the subfamily based on molecular and morphological data remains relatively poorly understood. Nevertheless, several relatively well-established relationships relevant to comparative bracovirus genomics are clear: a very close relationship between Cotesia Cameron and Glyptapanteles Ashmead, a cluster of related genera centered around Apanteles Foerster, and a relatively early divergence between Microplitis Foerster and its close relatives, and the other genera. Thus, we would expect the bracoviruses of Cotesia and Glyptapanteles to be relatively similar, and those of Microplitis to be among the most distant among the Microgastrinae.

**C. Bracovirus**

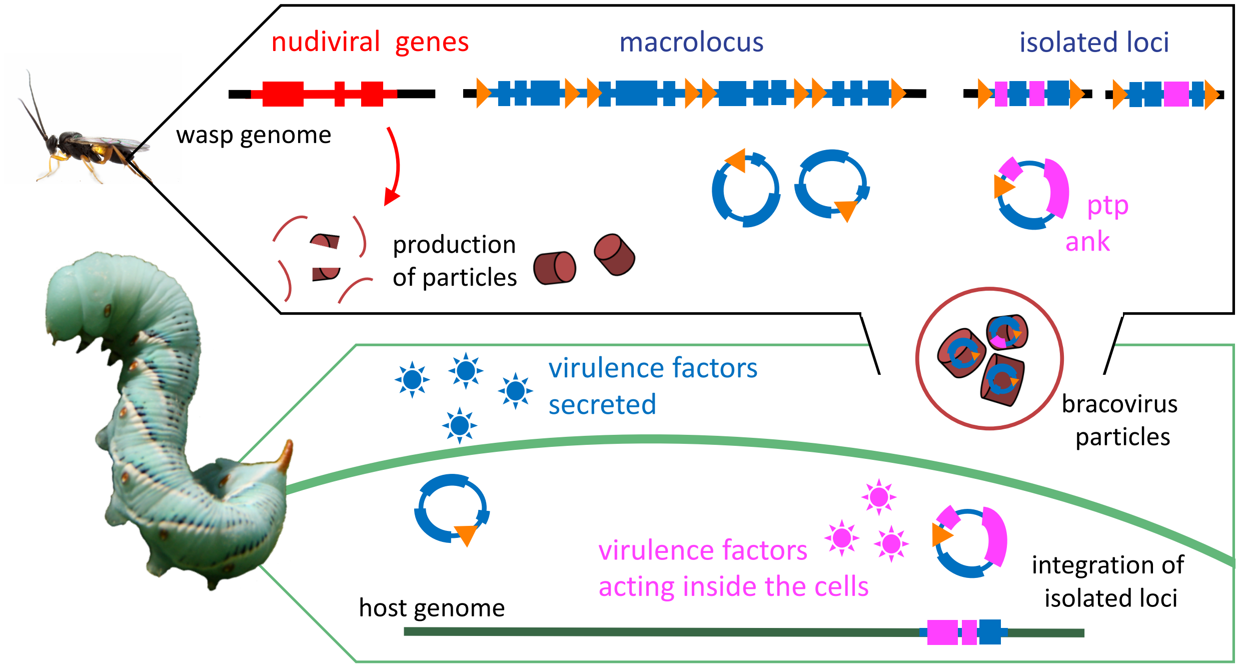

**Supplementary Figure 7.** Schematic representation of the bracovirus production in wasp ovaries and their function in host cells. Circles from isolated loci, encoding in particular *ptp* and *Vank* genes, integrate into parasitized host DNA using HIM site mediated mechanism.

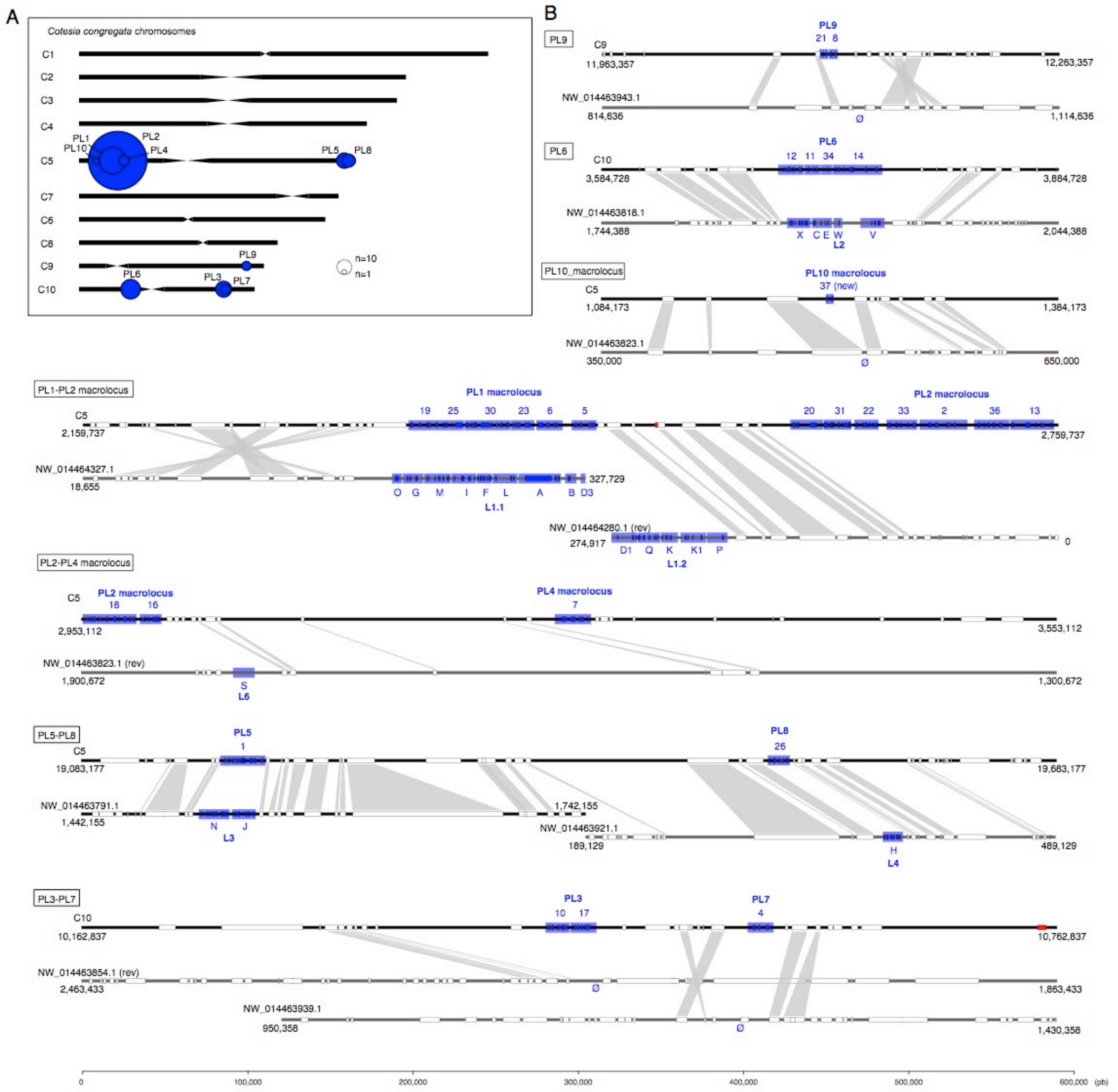

**Supplementary Figure 8. Synteny between Proviral Loci (PL) of *C. congregata*_and *M. demolitor*. A** *C. congregata* chromosomes map. **B** Comparison of *C. congregata* and *M. demolitor proviral loci*. *C. congregata* chromosomes and *M. demolitor* genome scaffolds are represented in black and dark grey respectively. Numbers 1 to 37 correspond to the 37 segments identified in *C. congregata* with the corresponding proviral loci indicating above and identified in *C. congregata* chromosomes map. Blue boxes indicate virulence genes while white boxes refer to non-virulence genes. *M. demolitor* scaffolds for which the orientation if reversed compared to *C. congregata* chromosomes are indicated by “rev”. The scale shows length in bp. Ø indicates the absence of orthologous segment in *M. demolitor* genome.

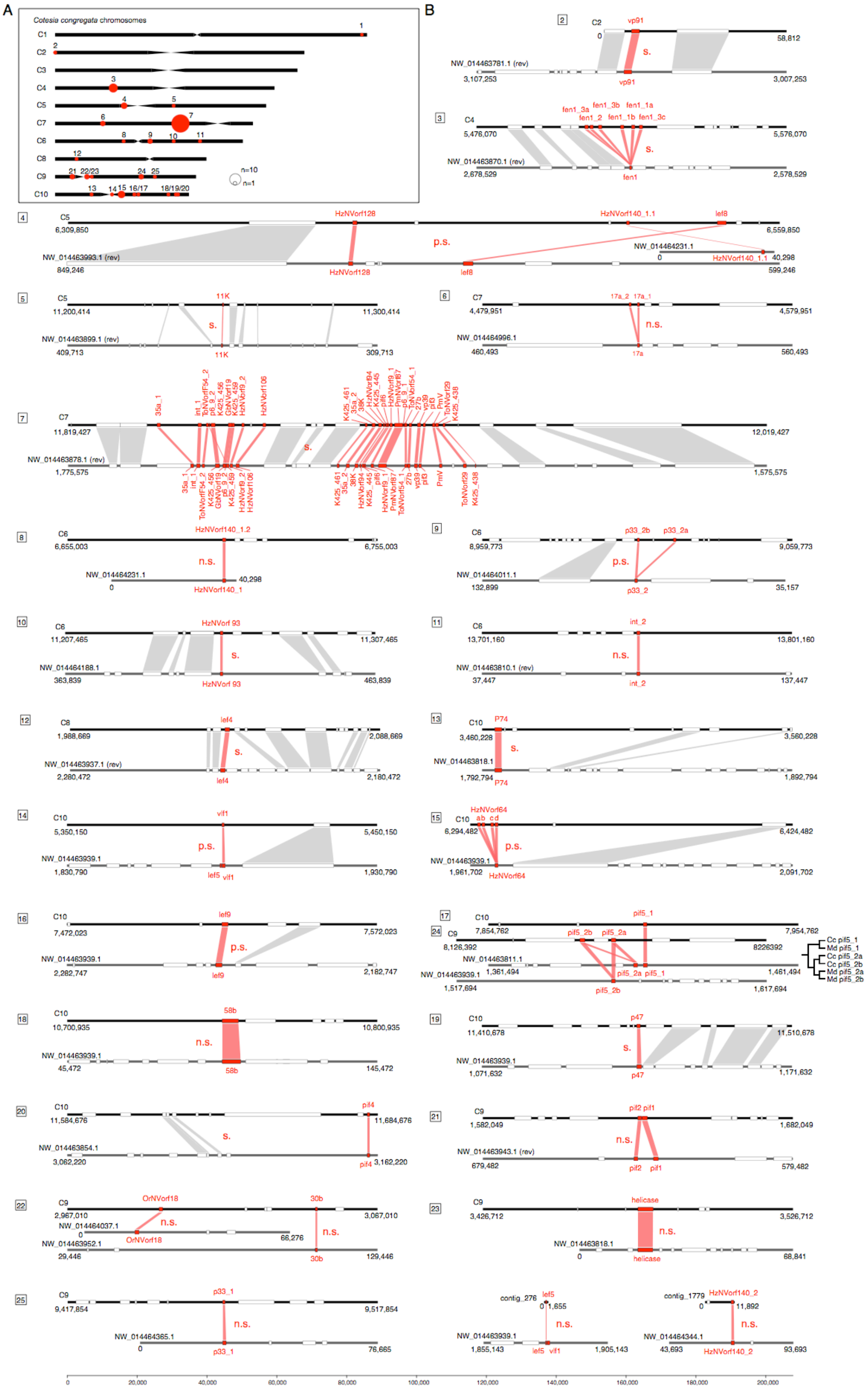

A
B
Cotesia congregata chromosomes
C1
C2
C3
C4
C5
C7
C6
C8
C9
C10
n=10
n=1

**Supplementary Figure 9. Synteny between nudiral genes containing regions of *C. congregata* and *M. demolitor*.** **A** *C. congregata* chromosomes map. **B** Comparison of nudiviral genes regions of *C. congregata* and *M. demolitor*. To validate a synteny between the two specie (indicated by "s." for synteny), we searched for at least two hymenopteran (non-viral) orthologous gene in the vicinity of homologous nudiviral gene(s) of the two species. If only one non-nudiviral orthologous gene was present we considered synteny as probable ("p.s.": probable synteny). Finally, when no orthologous gene was present in the vicinity of the nudiviral gene(s) in the two species, we considered the regions containing nudiviral genes were not homologous ("n.s": no synteny). *C. congregata* chromosomes and *M. demolitor* genome scaffolds are represented in black and dark grey respectively. Numbers 1 to 26 correspond to the 26 nudiviral loci identified in *C. congregata* chromosomes map. Red boxes indicate nudiviral genes and white boxes refer to hymenopteran genes. *M. demolitor* scaffolds for which the orientation is reversed compared to *C. congregata* chromosomes are indicated by "rev". The scale shows length in bp.

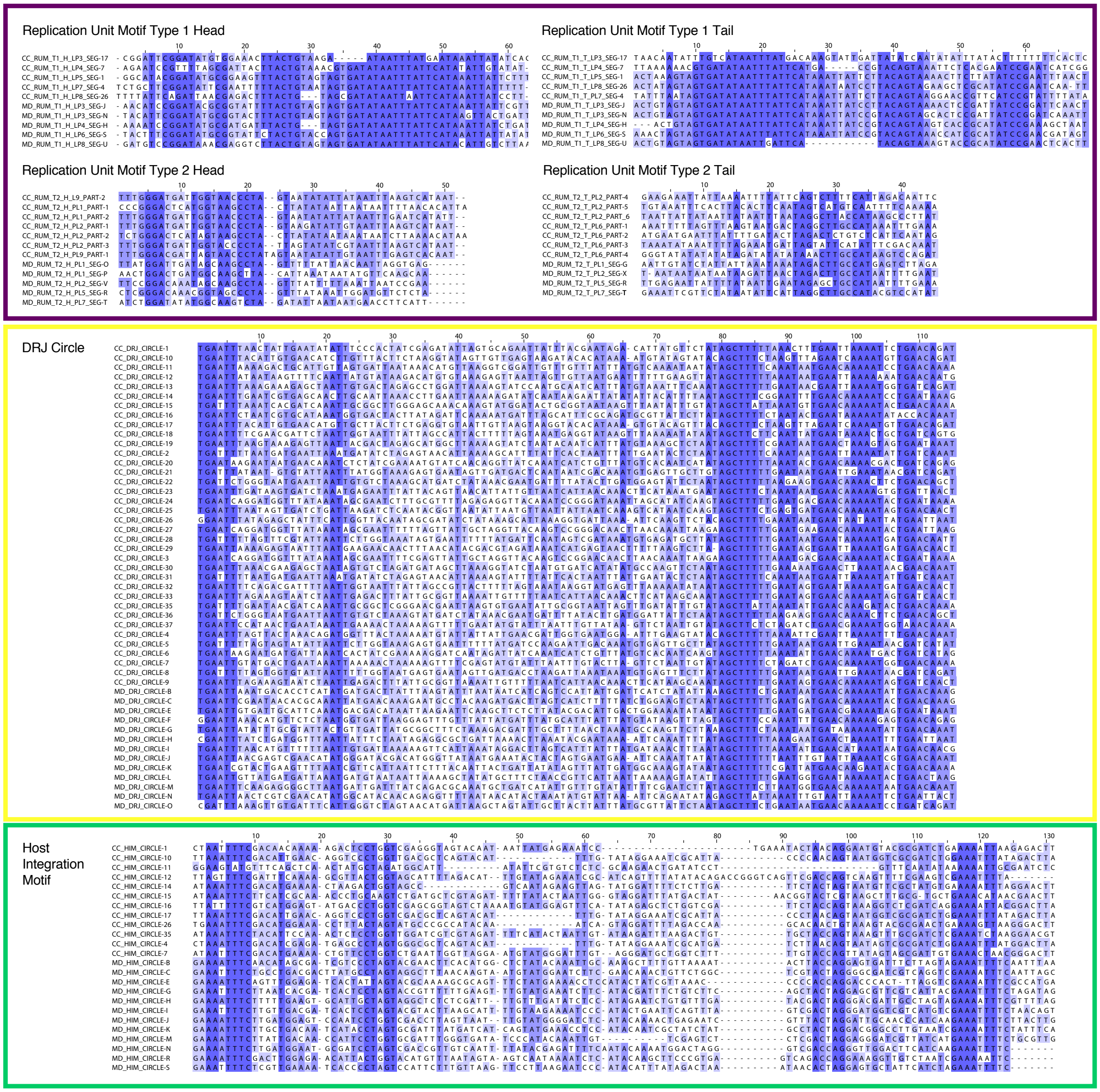

**Supplementary Figure 10**. In purple framework, the alignment of Replication Unit Motifs (RUM) including *C. congregata* and *M. demolitor* sequences. In yellow framework, the alignment of the circle Direct Repeat Junctions (DRJ). In green framework, the alignment of the Host Integration Motifs (HIM).

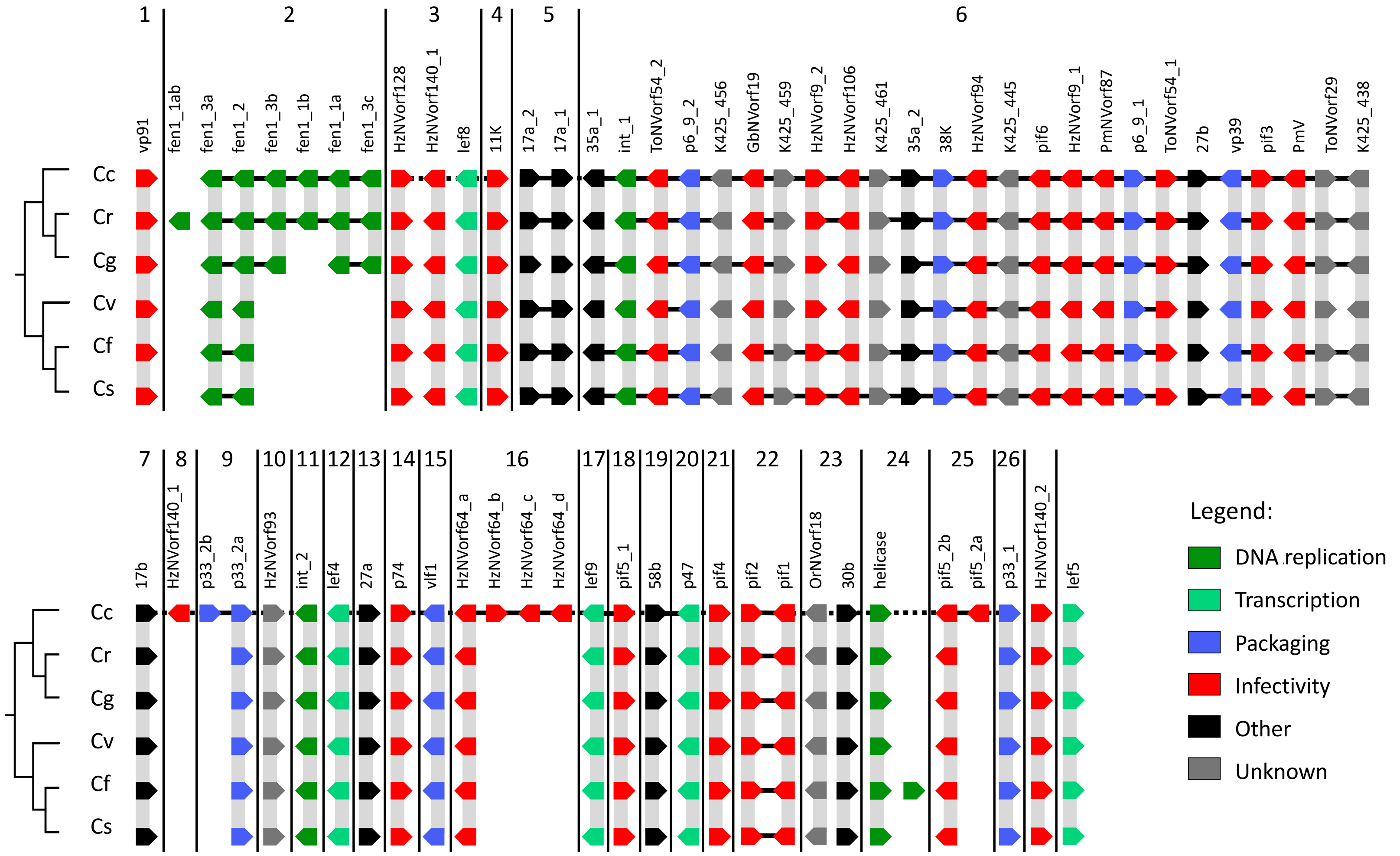

**Supplementary Figure 11**. Synteny of nudiviral genes across *Cotesia* species. Continuous black lines represent scaffolds and arrows indicate the orientation of the genes in each species. Number 1 to 26 correspond to the 26 nudiviral loci identified in *C. congregata* chromosomes. Cc, Cr, Cg, Cv, Cf, and Cs refer respectively to *C. congregata*, *C. rubecula*, *C. glomerata*, *C. vestalis*, *C. flavipes*, and *C. sesamiae*.

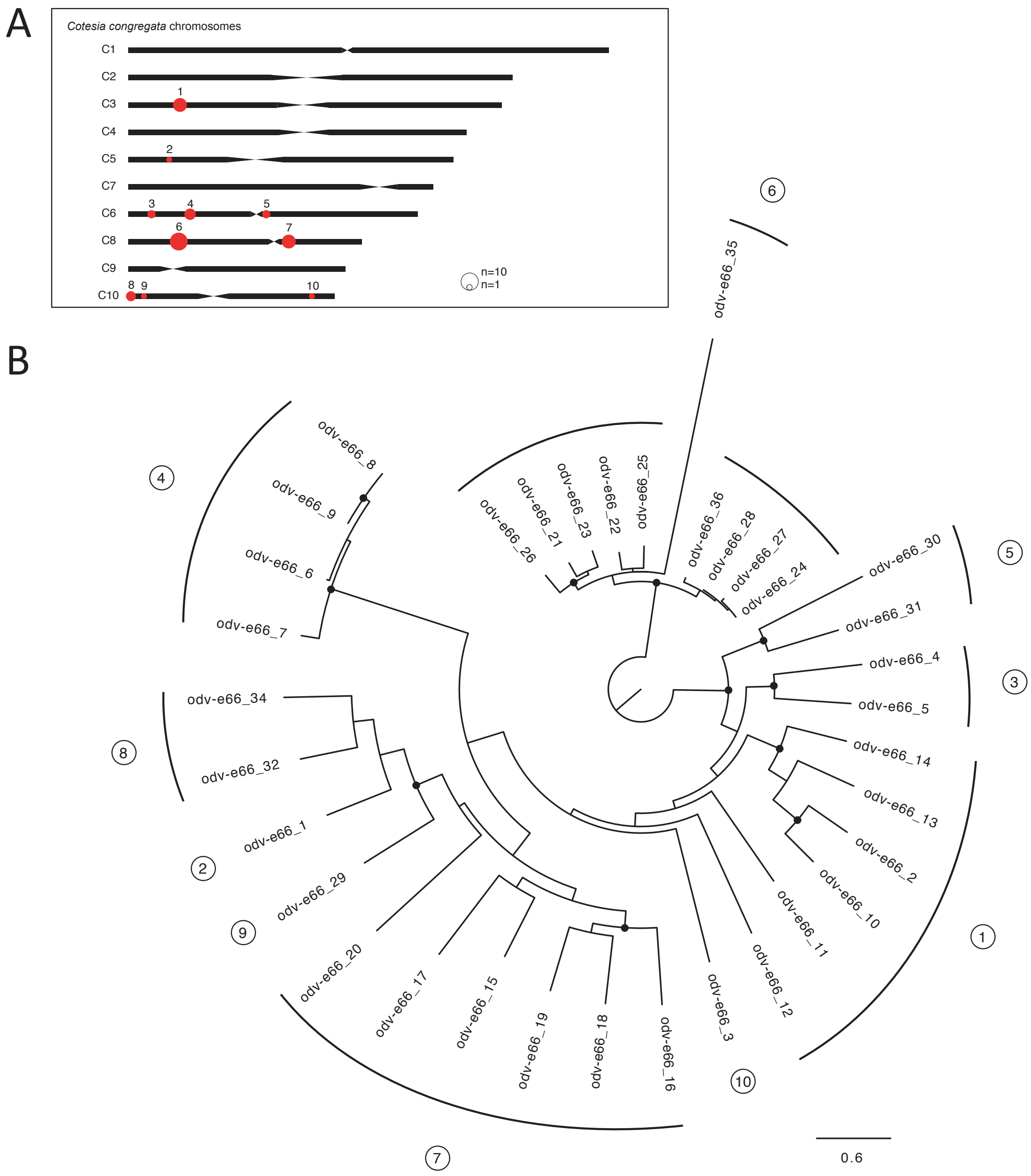

**Supplementary Figure 12.** Evolution of the *odv-e66* nudiviral gene family. **A.** Localization of odv-e66 genes on *C. congregata* chromosomes. The 36 *odv-e66* genes are clustered in 10 groups distributed in five different chromosomes of *C. congregata*. **B.** Maximum-likelihood phylogeny of *C. congregata* odv-e66 family (1,000 bootstraps). Prior tree construction, the regions that were present in less than 50% of the aligned sequences were manually curated from the codon-based alignment and the *odv-e66_33* gene was excluded due to its short sequence. The tree was rooted to the midpoint and the black dots indicate nodes with at least 80% of support.

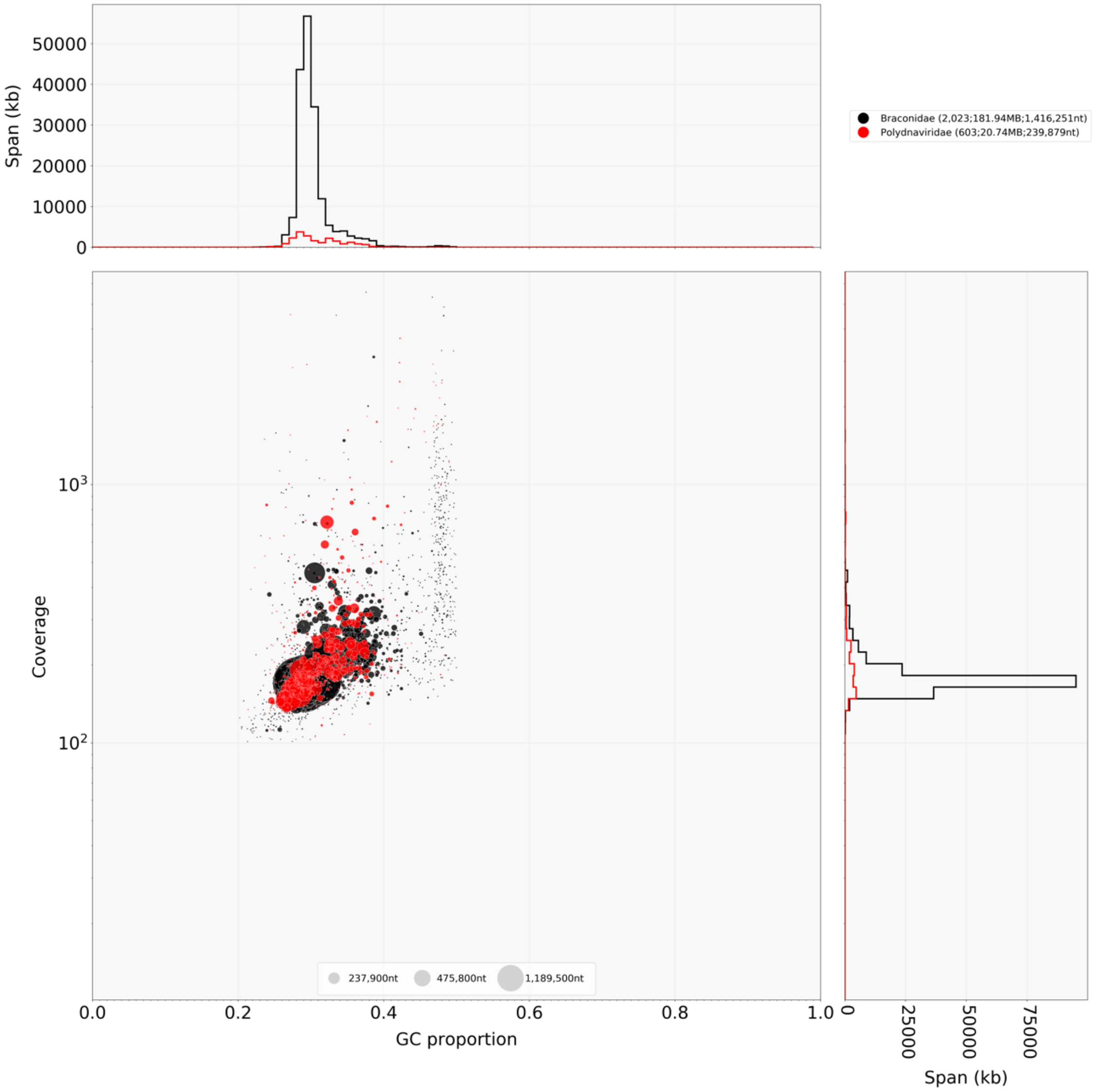

**Supplementary Figure 13.** Blob-plot or taxon-annotated GC content-coverage plot of *C. congregata* scaffolds. Each circle represents a scaffold in the assembly, scaled by length, and colored by order-level NCBI taxonomy assigned by BlobTools. The X axis corresponds to the average GC content of each scaffold and the Y axis corresponds to the average coverage based on alignment of Illumina reads. Marginal histograms show cumulative genome content (in Kb) for bins of coverage (Y axis) and GC content (X axis).

**Supplementary Table 8.** Evolutionary rates of nudiviral genes in *Cotesia* species (*C.c* = *C. congregata*; *C.r* = *C. rubecula*; *C.g* = *C. glomerata*; *C.v* = *C. vestalis*; *C.f* = *C. flavipes*; *C.s* = *C. sesamiae*; *M.d* = *M. demolitor.*

| Protein fonction | Gene name | Gene content by species | | | | | | | dN/dS value | p-value |
|---|---|---|---|---|---|---|---|---|---|---|
| | | *C.c* | *C. r* | *C. g* | *C. v* | *C. f* | *C. s* | *M. d* | | |
| Replication, DNA processing | *helicase* | + | + | + | + | + | + | + | 0.1343 | < 0.001 |
| | *int_1* | + | + | + | + | + | + | + | 0.15141 | < 0.001 |
| | *int_2* | + | + | + | + | + | + | + | 0.13921 | < 0.001 |
| | *fen-1-1a* | + | 2 | + | - | - | - | - | 0.75876 | 0.284 |
| | *fen-1-1b* | + | + | - | - | - | - | - | n.c. | n.c. |
| | *fen-1-2* | + | + | + | + | + | + | + | 0.27467 | < 0.001 |
| | *fen-1-3a* | + | + | + | + | + | + | - | 0.13039 | < 0.001 |
| | *fen-1-3b* | + | + | + | - | - | - | - | 0.87961 | 0.555 |
| | *fen-1-3c* | + | + | + | - | - | - | - | 0.38875 | < 0.001 |
| Transcription | *p47* | + | + | + | + | + | + | + | 0.05605 | < 0.001 |
| | *lef-8* | + | + | + | + | + | + | + | 0.05973 | < 0.001 |
| | *lef-9* | + | + | + | + | + | + | + | 0.05088 | < 0.001 |
| | *lef-4* | + | + | + | + | + | + | + | 0.08525 | < 0.001 |
| | *lef-5* | + | + | + | + | + | + | + | 0.36112 | < 0.001 |
| Packaging, assembly and release | *vlf-1* | + | + | + | + | + | + | + | 0.17119 | < 0.001 |
| | *vp91* | + | + | + | + | + | + | + | 0.3808 | < 0.001 |
| | *vp39* | + | + | + | + | + | + | + | 0.72163 | 0.057 |
| | *p33_1* | + | + | + | + | + | + | + | 0.17862 | < 0.001 |
| | *p33_2* | 2 | 1 | 1 | 1 | 1 | 1 | 1 | 0.15432* | < 0.001 |
| | *38K* | + | + | + | + | + | + | + | 0.19847 | < 0.001 |
| | *p6.9_1* | + | + | + | + | + | + | + | 0.08337 | < 0.001 |

| | | | | | | | | | | |
|---|---|---|---|---|---|---|---|---|---|---|
| | *p6.9_2* | + | + | + | + | + | + | + | 0.70662 | 0.267 |
| per os infectivity factors and ODV envelope particle components | *p74* | + | + | + | + | + | + | + | 0.79274 | 0.058 |
| | *pif-1* | + | + | + | + | + | + | + | 0.45494 | < 0.001 |
| | *pif-2* | + | + | + | + | + | + | + | 0.319 | < 0.001 |
| | *pif-3* | + | + | + | + | + | + | + | 0.71855 | 0.098 |
| | *pif-4* | + | + | + | + | + | + | + | 0.45842 | < 0.001 |
| | *pif-5_1* | + | + | + | + | + | + | + | 1.10226 | 0.38 |
| | *pif-5_2* | 2 | 1 | 1 | 1 | 1 | 1 | 3 | 0.3709* | < 0.001 |
| | *pif-6* | + | + | + | + | + | + | + | 0.41774 | < 0.01 |
| | *HzNVorf9_1* | + | + | + | + | + | + | + | 0.08198 | < 0.001 |
| | *HzNVorf9_2* | + | + | + | + | + | + | + | 0.08558 | < 0.001 |
| | *GbNVorf19* | + | + | + | + | + | + | + | 0.30979 | < 0.001 |
| | *HzNVorf64* | 4 | 1 | 1 | 1 | 1 | 1 | 1 | 0.09716* | < 0.001 |
| | *HzNVorf94* | + | + | + | + | + | + | + | 0.11672 | < 0.001 |
| | *HzNVorf106* | + | + | + | + | + | + | + | 0.08689 | < 0.001 |
| | *PmV* | + | + | + | + | + | + | + | 0.11206 | < 0.001 |
| | *11K* | + | + | + | + | + | + | + | 0.29473 | < 0.001 |
| | *HzNVorf128* | + | + | + | + | + | + | + | 0.11851 | < 0.001 |
| | *HzNVorf140_1* | 3 | 1 | 1 | 1 | 1 | 1 | 1 | 0.19362* | < 0.001 |
| | *HzNVorf140_2* | + | + | + | + | + | + | + | 0.1557 | < 0.001 |
| | *PmNVorf87* | + | + | + | + | + | + | + | 0.34099 | < 0.001 |
| | *ToNVorf54_1* | + | + | + | + | + | + | + | 0.245 | < 0.001 |
| | *ToNVorf54_2* | + | + | + | + | + | + | + | 0.23574 | < 0.001 |
| Other particle components | *17a_1* | + | + | + | + | + | + | + | 0.96093 | 0.839 |
| | *17a_2* | + | + | + | + | + | + | - | 0.62831 | < 0.05 |
| | *17b* | + | + | + | + | + | + | + | 0.11694 | < 0.001 |
| | *27a* | + | + | + | + | + | + | + | 0.2348 | < 0.001 |
| | *27b* | + | + | + | + | + | + | + | 0.24099 | < 0.001 |

| | | | | | | | | | | |
|---|---|---|---|---|---|---|---|---|---|---|
| | *30b* | + | + | + | + | + | + | + | 0.46248 | < 0.001 |
| | *35a_1* | + | + | + | + | + | + | + | 0.74071 | 0.088 |
| | *35a_2* | + | + | + | + | + | + | + | 0.55692 | < 0.001 |
| | *58b* | + | + | + | + | + | + | + | 0.58867 | < 0.001 |
| Unknown | *K425_438* | + | + | + | + | + | + | + | 0.66340 | 0.076 |
| | *K425_445* | + | + | + | + | + | + | + | 0.83681 | 0.45 |
| | *K425_456* | + | + | + | + | + | + | + | 0.14541 | < 0.001 |
| | *K425_459* | + | + | + | + | + | + | + | 0.20317 | < 0.001 |
| | *K425_461* | + | + | + | + | + | + | + | 0.14772 | < 0.001 |
| | *ToNVorf29* | + | + | + | + | + | + | + | 0.15643 | < 0.001 |
| | *HzNVorf93* | + | + | + | + | + | + | + | 0.18237 | < 0.001 |
| | *OrNVorf18* | + | + | + | + | + | + | + | 0.36808 | < 0.001 |
| *odv-e66* | *odv-e66-1* | + | + | + | + | + | + | - | 0.57915 | < 0.001 |
| | *odv-e66-2* | + | + | + | + | + | + | + | 0.84754 | < 0.001 |
| | *odv-e66-3* | + | + | + | + | + | + | + | 0.51583 | < 0.01 |
| | *odv-e66-4* | + | + | + | + | + | + | - | 0.53930 | < 0.001 |
| | *odv-e66-5* | + | + | + | + | + | + | - | 0.71140 | < 0.05 |
| | *odv-e66-6/7/8/9* | 4 | 1 | 1 | 3 | 1 | 1 | 2 | n.c. | n.c. |
| | *odv-e66-10* | + | + | - | + | + | + | - | 1.10867 | 0.589 |
| | *odv-e66-11* | + | + | + | + | + | + | - | 0.25798 | < 0.001 |
| | *odv-e66-12* | + | + | + | + | + | + | - | 0.23769 | < 0.001 |
| | *odv-e66-13* | + | + | + | + | + | + | + | 0.87336 | 0.439 |
| | *odv-e66-14* | + | + | + | + | + | + | + | 0.61298 | < 0.01 |
| | *odv-e66-15* | + | + | + | + | + | + | + | 0.25491 | < 0.001 |
| | *odv-e66-16* | + | + | + | + | + | + | + | 0.51120 | < 0.001 |
| | *odv-e66-17* | + | + | + | + | + | + | + | 0.58245 | < 0.001 |
| | *odv-e66-18* | + | + | + | + | + | + | - | 0.41734 | < 0.001 |

| | | | | | | | | | | |
|---|---|---|---|---|---|---|---|---|---|---|
| | *odv-e66-19* | + | + | + | + | + | + | + | 0.49926 | < 0.001 |
| | *odv-e66-20* | + | + | + | + | + | + | + | 0.17390 | < 0.001 |
| | *odv-e66-21/22/23/ 24/25/26/27/28* | 8 | 1 | - | - | - | - | - | n.c. | n.c. |
| | *odv-e66-29* | + | + | + | + | + | + | 2 | 0.36489 | < 0.001 |
| | *odv-e66-30* | + | + | + | + | + | + | - | 0.85965 | 0.211 |
| | *odv-e66-31* | + | + | - | - | - | - | - | n.c. | n.c. |
| | *odv-e66-32* | + | + | - | + | + | + | - | 0.52025 | < 0.001 |
| | *odv-e66-33* | + | + | - | - | - | - | - | n.c. | n.c. |
| | *odv-e66-34* | + | + | - | - | - | - | - | n.c. | n.c. |
| | *odv-e66-35* | + | - | - | - | - | - | - | n.c. | n.c. |
| | *odv-e66-36* | + | - | - | - | - | - | - | n.c. | n.c. |

* the closest orthologue among *C.congregata* duplications was used to estimate dN/dS
n.c. – dN/dS was not etimated when orthologues and paralogues were not distinguishable

#### D. Immunity

**Supplementary Table 9.** Immunity gene repertoires of Hymenoptera. The table indicates the number of genes annotated (including incomplete genes and pseudogenes) in six hymenopteran species.

| Gene function | Pathway | Gene | Hymenoptera | | | Diptera | Lepidoptera | Hemiptera |
|---|---|---|---|---|---|---|---|---|
| | | | *C. congregata* | *N. vitripenis* | *A. mellifera* | *D. melanogaster* | *M. sexta* | *A. pisum* |
| Recognition | | PGRP | 6 | 11 | 4 | 13 | 14 | 0 |
| | | GNBP/βGBP | 2 | 3 | 2 | 3 | 5 | 2 |
| | | C type lectin | 2 | 28 | 12 | 34 | 34 | 5 |
| | | Hemolectin | 1 | 0 | 1 | 1 | 2 | 0 |
| | | Galectin | 3 | 3 | 2 | 6 | 4 | 2 |
| | | TEP | 3 | 3 | 4 | 6 | 3 | 2 |
| | | Dscam | 5 | 1 | 1 | 1 | 1 | 5 |
| | | Integrin | 7 | 0 | 0 | 2 | 4 | 7 |
| Signal transduction | | Serine protease | 58 | 55 | 57 | 12 | 107 | 36 |
| | | CLIP serine protease | 8 | 28 | 18 | 45 | 54 | 3 |
| | | Serpins | 4 | 12 | 7 | 29 | 34 | 14 |
| | Toll | Spätzle | 3 | 6 | 2 | 6 | 8 | 10 |
| | | Toll receptor | 7 | 6 | 5 | 9 | 16 | 7 |
| | | MyD88 | 1 | 1 | 1 | 1 | 1 | 1 |
| | | Tube | 2 | 1 | 1 | 1 | 1 | 1 |
| | | Pelle | 2 | 1 | 1 | 1 | 1 | 1 |

| | | | | | | | | |
|---|---|---|---|---|---|---|---|---|
| | | ANK/Cactus | 1 | 1 | 3 | 1 | 1 | 1 |
| | | Dif/Dorsal | 1 | 2 | 2 | 2 | 5 | 2 |
| Signalling pathways | IMD/JNK | IMD | 1 | 1 | 1 | 1 | 1 | 0 |
| | | FADD | 1 | n.d. | 1 | 1 | 1 | 1 |
| | | Dredd_IMD | 1 | 1 | 1 | 1 | 1 | 0 |
| | | Tab2_IMD | 1 | 1 | 2 | 1 | 1 | 1 |
| | | IAP2 | 5 | n.d. | 1 | 1 | 1 | 1 |
| | | ird5-IKKg | 1 | 1 | 1 | 1 | 1 | 1 |
| | | TAK1_IMD | 1 | n.d. | 1 | 1 | 1 | 1 |
| | | IKKg (Kenny) | 0 | 1 | 1 | 1 | 1 | 0 |
| | | Bendless-Ubc13 | 1 | n.d. | 1 | 1 | 1 | 1 |
| | | Relish_IMD | 1 | 3 | 1 | 1 | 1 | 0 |
| | | JNK kinase (Hemipterous-MKK7) | 1 | n.d. | 1 | 3 | 1 | 1 |
| | | Basket-JNK | 1 | 1 | 1 | 1 | 1 | 1 |
| | | Fos - kayak transcription factor | 1 | 1 | 1 | 1 | 1 | 0 |
| | | c-Jun | 1 | n.d. | 1 | 1 | 1 | 1 |
| | JAK/STAT | Upd3 | 0 | n.d. | 0 | 1 | 0 | 0 |
| | | PIAS | 3 | 1 | 3 | 1 | 1 | n.d. |
| | | SOCS | 1 | n.d. | 2 | 1 | 1 | 3 |
| | | Domeless | 1 | 1 | 1 | 1 | 1 | 3 |
| | | Hopscotch (JAK) | 1 | 1 | 4 | 1 | 1 | 1 |
| | | STAT | 1 | n.d. | 1 | 1 | 1 | 2 |
| | Melanization | Pro Phenol Oxidase | 1 | 3 | 1 | 3 | 2 | 2 |
| | Anti-microbial peptides | Hymenoptaecin | 1 | 1 | 1 | 0 | 0 | 0 |
| | | Moricin | 0 | 0 | 0 | 0 | 6 | 0 |
| | | Lebocin | 0 | 0 | 0 | 0 | 4 | 0 |
| | | Defensin | 6 | 2 | 2 | 1 | 24 | 0 |

| | | | | | | | | |
|---|---|---|---|---|---|---|---|---|
| | | Lysozyme | 1 | 1 | 3 | 13 | 6 | 3 |
| Effectors | | Chitinase | 11 | 4 | 5 | 16 | 11 | 7 |
| | | Transferrin | 3 | 2 | 1 | 2 | 4 | 2 |
| | | Peroxidase | 14 | 15 | 13 | 20 | 3 | 15 |
| | Others | Nitric Oxide Synthase (NOS) | 1 | 1 | 1 | 1 | 2 | 1 |
| | | Super Oxide Dismutase | 4 | 4 | 2 | 4 | 4 | 4 |
| | | Glutathione S Transferase | 8 | 7 | 5 | 35 | 31 | 18 |
| | | MIF | 0 | 0 | 0 | 0 | 1 | 5 |
| | | Heat Shock Protein | 11 | 4 | 4 | 13 | 16 | 15 |
| Antiviral immuniy | RNAi pathways | Dicer 1 | 1 | 0 | 1 | 1 | 1 | 1 |
| | | Dicer 2 | 1 | 0 | 1 | 1 | 1 | 1 |
| | | Argonaute | 3 | 0 | 1 | 3 | 1 | 4 |
| | | R2D2 | 1 | 0 | 1 | 1 | 1 | 1 |
| | | Vago | 0 | 0 | 1 | 1 | 1 | 0 |

n.d. for not determined

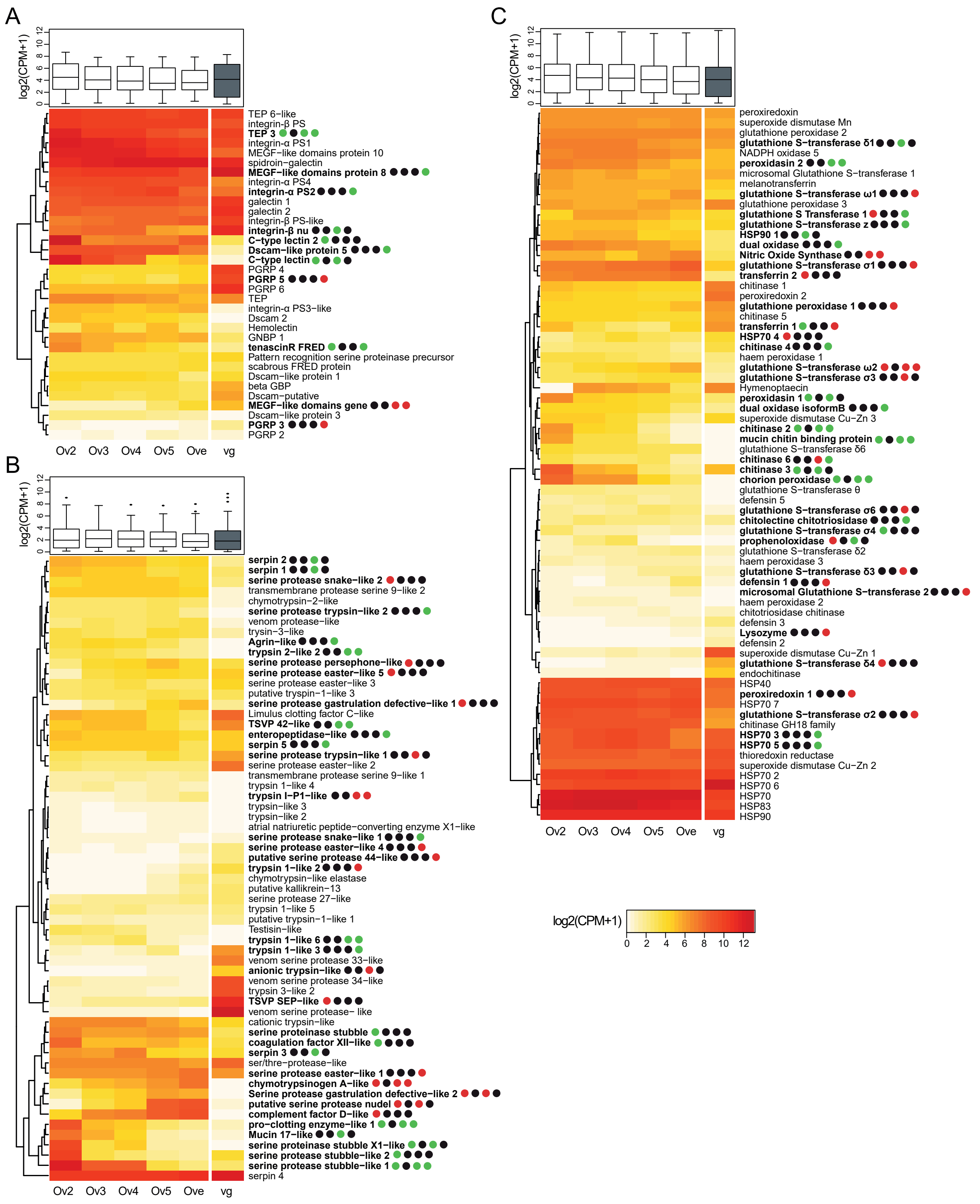

**Supplementary Figure 14.** Gene expression of immunity genes during *Cotesia congregata* development. Heatmaps show the expression of the genes involved in **A** signal recognition, **B** signal transduction and **C** effector functions across the developmental stages of ovaries (Ov2, Ov3, Ov4, Ov5, Ove) and in venom glands (vg). The trees on the left are unsupervised hierarchical clustering of expression values. Boxplots represent overall expression of each immunity gene group in ovaries and venom glands. Bold names highlight the genes that are differentially expressed and dots represent the four different comparisons studied between consecutive ovary stages (Ov2 *vs.* Ov3, Ov3 *vs*. Ov4, Ov4 *vs*. Ov5 and Ov5 *vs.* Ove). Black, red and green dots indicate similar, increased and reduced expressions between consecutive developmental stages respectively.

**E. Chemoreceptor**

To find their hosts, female wasps follow scents emitted by caterpillars and the plants they damage. The host identification process for oviposition acceptance occurs mainly during contact between the parasitoid and its host, when host products related to feeding activities, fecal pellets and oral secretions, play a crucial role. In insects, chemical signals are detected by sensory neurons expressing transmembrane receptor genes belonging to three different families: the odorant receptors (ORs) that are devoted to olfaction, the gustatory receptors (GRs) which are involved in taste, and the ionotropic receptors (IRs) that include receptors used in both chemosensory modalities[18]. We annotated these chemoreceptor gene families in the genome of *C. congregata*, and identified genes encoding 243 ORs and 54 GRs. These numbers are in the range of those obtained for other parasitoid wasps, either within the family Braconidae or in *N. vitripennis*, but remain lower than in ants (Supplementary Table 10). We also identified 105 IRs, whereas only 56 have been annotated in *Diachasma alloeum*, another Braconidae.

Focusing on ORs, we performed expert annotation and manual curation, resulting in 197 full-length gene sequences (encoding >350 amino acids) and 46 incomplete genes. A phylogenetic analysis of ORs from *C. congregata* and four other Hymenoptera species (*M. demolitor*, *N. vitripennis*, *A. mellifera*) showed that *C. congregata* ORs belong to 15 of the 18 strongly supported monophyletic OR lineages (aLRT ≥ 0.9) (Fig. 2), which have been previously described in Apocrita[30]. Using this phylogeny, we analyzed the dynamics of OR gene gains and losses and found that the equally high number of OR genes in *N. vitripennis* and in Braconidae (*M. demolitor* and *C. congregata*) results from independent expansions (Supplementary Fig.14). The number of OR genes in the last common ancestor of these parasitoid wasps may have been rather low (~60), and many duplication events occurred after the split between Braconidae and other parasitoid wasps. The most spectacular Braconidae-specific expansions occurred in clades B, C/D/E, F, Q/R/S and 9-exon, each harboring at least 25 genes in *C. congregata* (Fig 2; Supplementary Table 11). As expected, highly duplicated OR genes in *C. congregata* were found in 6 clusters of at least 10 tandemly arrayed genes, the largest one containing 19 genes (Supplementary Fig. 15). Even within Braconidae, many duplications occurred in ancestors of *Cotesia* species after the split with the lineage of *M. demolitor* (Fig. 2). We also identified Braconidae-specific gene losses, notably in subfamilies A, V and Z. This illustrates how dynamic is the evolution of OR genes within parasitoid wasps.

In an attempt to study whether different host specificities of *Cotesia* species could be linked with differences in OR repertoires, we annotated OR genes in four other *Cotesia* species. *C. congregata*, *C. sesamiae* and *C. flavipes* are parasitizing a large range of lepidopteran species, whereas *C. rubecula* and *C. vestalis* are specialists on *Pieris rapae* and *Plutella xylostella* larvae, respectively. Interestingly, OR copy numbers varied significantly during the evolution of the genus *Cotesia*, and higher numbers of OR genes were found in the two specialist species (Supplementary Table 11). The lack of phylogenetic resolution for closely related *Cotesia* OR genes precluded any comprehensive analysis of gene gains and losses, but we found several expansions in *C. rubecula* and *C. vestalis* within many OR clades (Supplementary Fig. 16).

**Supplementary Table 10.** Chemoreceptor gene repertoires of Hymenoptera. The table indicates the number of OR, GR and IR genes annotated (including incomplete genes and pseudogenes) in seven hymenopteran species.

| | Braconidae | | | Pteromalidae | Formicidae | Apidae | Cephidae |
|---|---|---|---|---|---|---|---|
| Species | *C. congregata* | *M. demolitor* | *D. alloeum* | *N. vitripennis* | *P. barbatus* | *A. mellifera* | *C. cinctus* |
| ORs | 243 | 203 | 201 | 216 | 399 | 162 | 73 |
| GRs | 54 | 86 | 40 | 58 | 73 | 13 | 35 |
| IRs | 105 | n.d. | 56 | 153 | 24 | 21 | 49 |
| Citation | This study | 26 | 27 | 28<br>29<br>30 | 31<br>30 | 27<br>28<br>30<br>32 | 28 |

n.d. for not determined

**Supplementary Table 11**. OR copy number per subfamily[26] compared to data obtained from genomes of *Cotesia* species.

| OR subfamily | *C. congregata* | *C. rubecula* | *C. vestalis* | *C. flavipes* | *C. sesamiae* | *M. demolitor* | *N. vitripennis* | *A. mellifera* | *C. cinctus* |
|---|---|---|---|---|---|---|---|---|---|
| A | 0 | 0 | 0 | 0 | 0 | 0 | 0 | 3 | 1 |
| B | 34 | 42 | 33 | 28 | 29 | 27 | 0 | 1 | 1 |
| CDE | 34 | 42 | 39 | 33 | 28 | 36 | 34 | 7 | 14 |
| F | 28 | 25 | 28 | 22 | 23 | 20 | 20 | 1 | 0 |
| GHX | 21 | 22 | 19 | 15 | 17 | 16 | 12 | 16 | 1 |
| J | 1 | 1 | 1 | 1 | 1 | 3 | 0 | 19 | 12 |
| KL | 16 | 20 | 12 | 9 | 9 | 14 | 8 | 59 | 8 |
| MNOP | 12 | 18 | 13 | 10 | 12 | 8 | 3 | 6 | 0 |
| QRS | 25 | 34 | 23 | 15 | 16 | 21 | 4 | 1 | 11 |
| TU | 11 | 10 | 12 | 9 | 9 | 12 | 22 | 3 | 4 |
| V | 0 | 0 | 0 | 0 | 0 | 0 | 9 | 6 | 1 |
| Z | 0 | 0 | 0 | 0 | 0 | 0 | 13 | 1 | 2 |
| ZB | 14 | 24 | 19 | 17 | 11 | 10 | 19 | 0 | 1 |
| 9-exon | 34 | 45 | 40 | 32 | 31 | 31 | 61 | 34 | 2 |
| Other | 13 | 13 | 13 | 11 | 11 | 5 | 11 | 5 | 15 |
| Total | 243 | 296 | 252 | 202 | 197 | 203 | 216 | 162 | 73 |

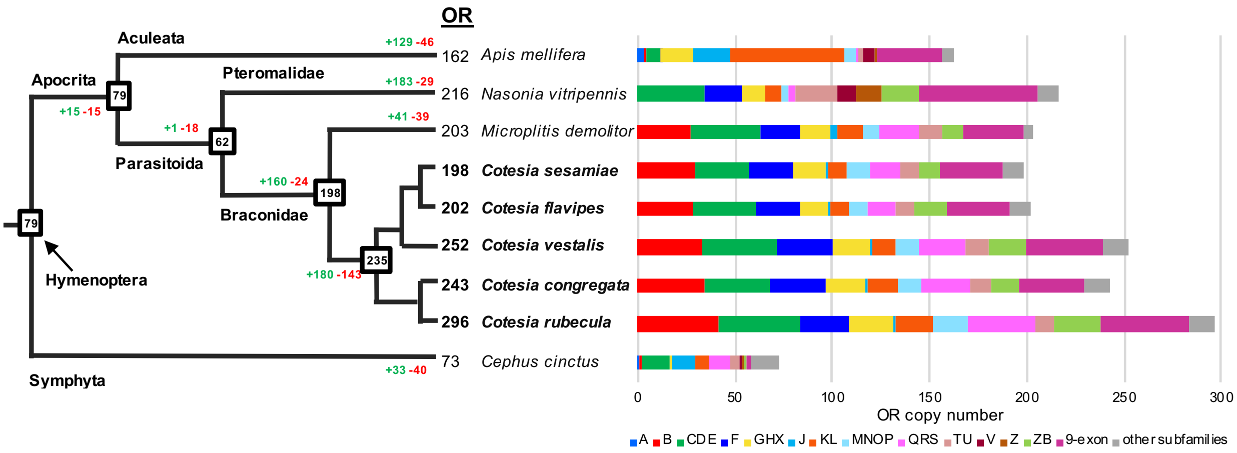

**Supplementary Figure 15.** Copy number dynamics of OR genes in five Cotesia species and four other Hymenoptera species. Estimated numbers of gene gain and loss events are shown on each branch of the species tree in green and red, respectively. The size of OR repertoires in common ancestors is indicated in boxes at the corresponding nodes of the species tree. The histogram represents the distribution of OR copy number per subfamily for each species.

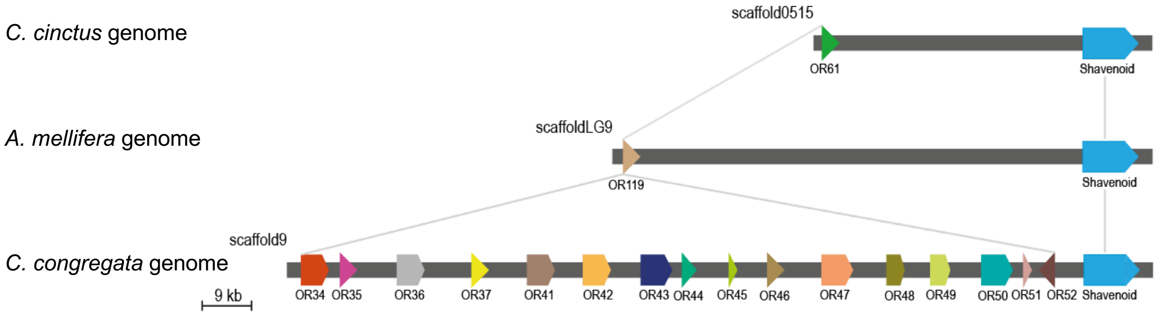

**Supplementary Figure 16.** Synteny among the OR subfamily B in *C. congregata*, *C. cinctus* and *A. mellifera*, based on inter-specific conservation of the shavenoid gene, showing expansion through tandem duplications for *C. congregate* OR genes. Orientation (arrows) of genes within scaffolds are indicated.

A

Subfamily QRS

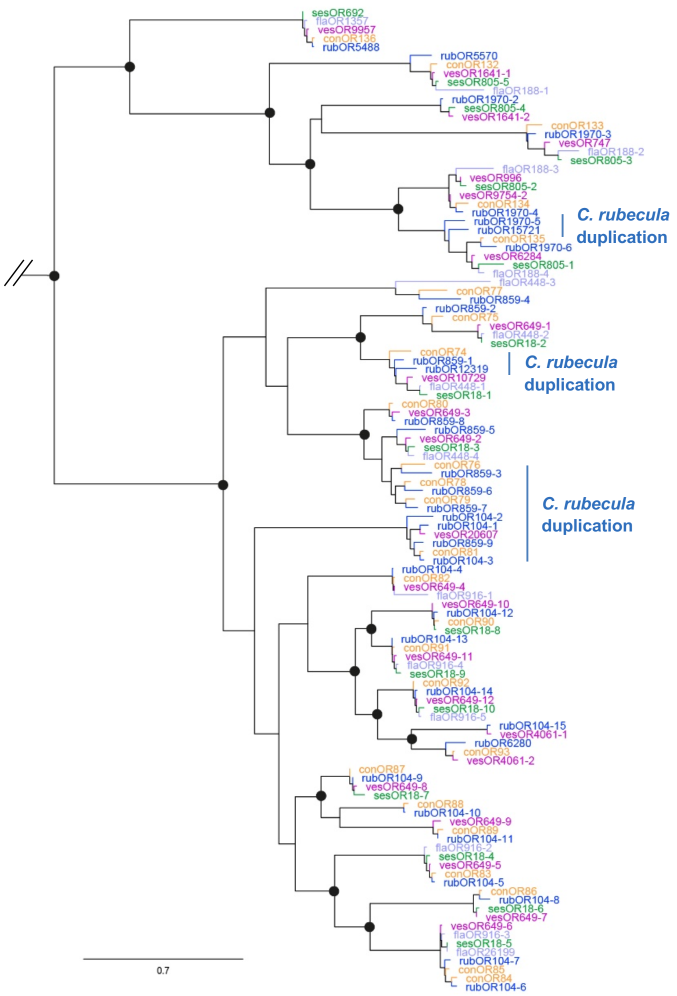

B

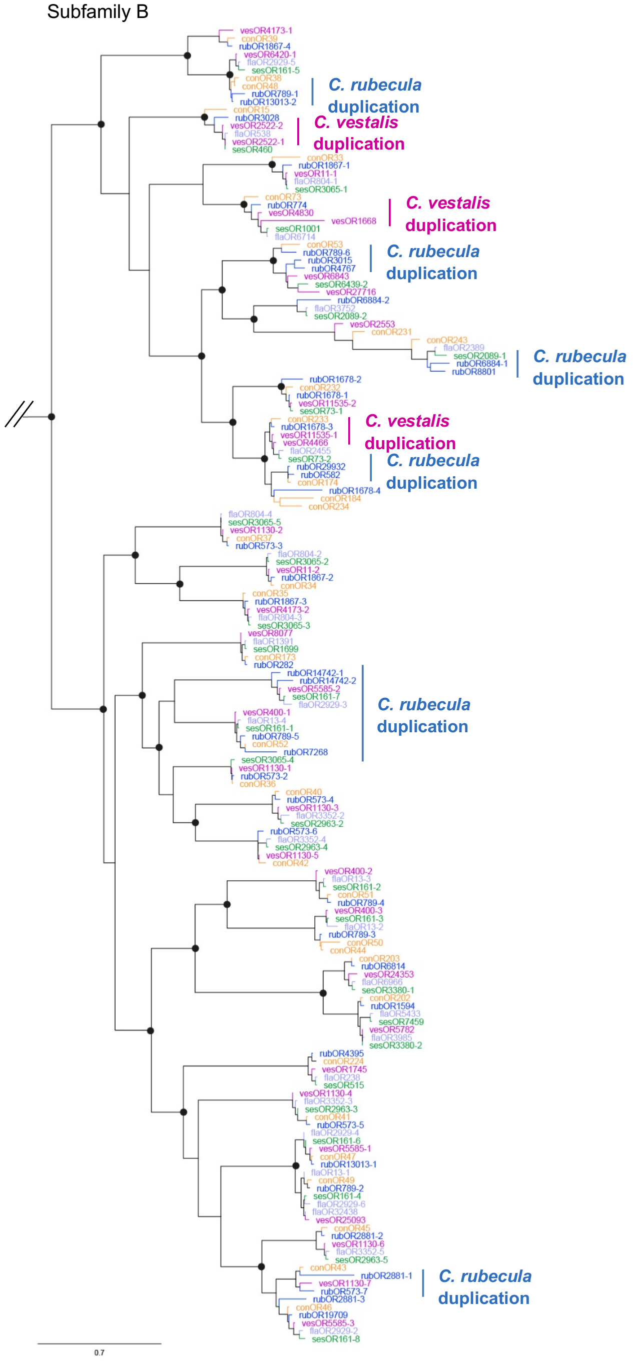

**Supplementary Figure 17.** Phylogenies of OR subfamilies **A** QRS and **B** B in five *Cotesia* species. These subtrees were extracted from maximum-likelihood phylogeny of *Cotesia* ORs including OR repertoires from *C. congregata* (con, orange), *C. flavipes* (fla, purple), *C. rubecula* (rub, blue), *C. sesamiae* (ses, green) and *C. vestalis* (ves, pink). The tree was rooted using the Orco clade as outgroup. Circles indicate nodes strongly supported by the approximate likelihood-ratio test (aLRT ≥ 0.95). The scale bar represents 0.7 expected amino acid substitutions per site. *C. rubecula* and *C. vestalis* duplications are highlighted on the right.

#### F. Detoxification

**Supplementary Table 12.** Detoxification gene repertoires of Hymenoptera. The table indicates the number of genes annotated (including incomplete genes and pseudogenes) in six hymenopteran species.

| | Hymenoptera | | | Diptera | Lepidoptera | Hemiptera |
|---|---|---|---|---|---|---|
| Enzyme family | *C. congregata* | *N. vitripennis* | *A. mellifera* | *D. melanogaster* | *M. sexta* | *A. pisum* |
| P450 | 70 | 87 | 46 | 86 | 103 | 64 |
| CCE | 35 | 41 | 24 | 35 | 96 | 30 |
| GST | 17 | 19 | 10 | 40 | 31 | 20 |
| UGT | 11 | 22 | 12 | 34 | 43 | 57 |
| ABC | 46 | 55 | 41 | 56 | 53 | 70 |
| Citation | This study | [33] | [34] [35] | [36] [37] | [38] [39] | [40] |

**Supplementary Table 13.** Detoxification gene repertoires of *Cotesia* species. The table indicates the number of genes annotated (including incomplete genes and pseudogenes) in six *Cotesia* species.

| Enzyme family | *C. congregata* | *C. rubecula* | *C. glomerata* | *C. vestalis* | *C. flavipes* | *C. sesamiae* |
|---|---|---|---|---|---|---|
| P450 | 70 | 68 | 70 | 65 | 48 | 50 |
| CCE | 26 | 32 | 30 | 27 | 22 | 24 |
| GST | 17 | 18 | n.d. | 17 | 17 | 17 |
| UGT | 10 | 10 | n.d. | 10 | 10 | 10 |
| ABC | 46 | 49 | 47 | 46 | 44 | 44 |

n.d. for not determined

**Supplementary Table 14.** Detailled detoxification gene repertoires of *Cotesia* species. The table indicates the number of genes annotated (including incomplete genes and pseudogenes) in six *Cotesia* species.

| | | *C. congregata* | *C. rubecula* | *C. glomerata* | *C. vestalis* | *C. flavipes* | *C. sesamiae* |
|---|---|---|---|---|---|---|---|
| P450 | CYP2 clan | 10 | 9 | n.d. | 9 | 9 | 9 |
| | CYP3 clan | 36 | 33 | n.d. | 31 | 25 | 24 |
| | CYP4 clan | 18 | 15 | n.d. | 14 | 9 | 10 |
| | mito clan | 6 | 6 | n.d. | 6 | 6 | 6 |
| CCE | Clade A | 14 | 16 | 12 | 14 | 12 | 13 |
| | Clade B | 5 | 8 | 10 | 8 | 5 | 6 |
| | Clade D | 1 | 2 | 2 | 1 | 1 | 1 |
| | Clade E | 4 | 4 | 4 | 2 | 2 | 2 |
| | Clade F | 2 | 2 | 2 | 2 | 2 | 2 |
| GST | Delta | 5 | 7 | n.d. | 6 | 6 | 6 |
| | Epsilon | 0 | 0 | n.d. | 0 | 0 | 0 |
| | Omega | 2 | 2 | n.d. | 2 | 2 | 2 |
| | Sigma | 5 | 5 | n.d. | 5 | 5 | 5 |
| | Theta | 1 | 1 | n.d. | 1 | 1 | 1 |
| | Zeta | 1 | 1 | n.d. | 1 | 1 | 1 |
| | microsomal | 2 | 16 | n.d. | 15 | 15 | 15 |
| | unclassified | 2 | 2 | n.d. | 2 | 2 | 2 |
| UGT | classe 1 | 0 | 0 | n.d. | 0 | 0 | 0 |
| | classe 2 | 1 | 1 | n.d. | 1 | 1 | 1 |
| | classe 3 | 0 | 0 | n.d. | 0 | 0 | 0 |
| | classe 4a | 4 | 4 | n.d. | 4 | 4 | 4 |
| | classe 4b | 1 | 1 | n.d. | 1 | 1 | 1 |
| | classe 4c | 2 | 2 | n.d. | 2 | 2 | 2 |
| | classe 5 | 1 | 1 | n.d. | 1 | 1 | 1 |
| | classe 6 | 1 | 1 | n.d. | 1 | 1 | 1 |
| | unclassified | 0 | 0 | n.d. | 0 | 0 | 0 |
| ABC | subfamily A | 4 | 4 | 4 | 4 | 4 | 4 |
| | subfamily B | 4 | 4 | 3 | 3 | 3 | 3 |
| | subfamily C | 16 | 17 | 17 | 14 | 16 | 16 |
| | subfamily D | 2 | 2 | 2 | 2 | 2 | 2 |
| | subfamily E | 1 | 2 | 1 | 2 | 1 | 0 |
| | subfamily F | 3 | 4 | 3 | 4 | 3 | 3 |

| | | | | | | | |
|---|---|---|---|---|---|---|---|
| | subfamily G | 13 | 13 | 14 | 14 | 12 | 13 |
| | subfamily H | 3 | 3 | 3 | 3 | 3 | 3 |

n.d. for not determined
